# End-to-end single-stranded DNA sequence design with all-atom structure reconstruction

**DOI:** 10.64898/2025.12.05.692525

**Authors:** Yunda Si, Yaochen Xu, Luonan Chen

## Abstract

Designing biological sequences that fold into predefined conformations is a central challenge in bioengineering. Although deep learning has enabled significant advances in protein and RNA sequence design, progress in single-stranded DNA (ssDNA) design has been constrained by the limited availability of structural data. To address this challenge, we introduce InvDNA, a deep learning-based method that designs ssDNA sequences directly from backbone atomic coordinates. This end-to-end formulation avoids the loss of structural information during backbone-to-feature conversion and further accommodates flexible backbone representations, dynamic sequence masking, and structural reconstruction objectives. These strategies bolster InvDNA’s ability to generalize across diverse ssDNA structural contexts while enabling additional functionalities, including generating diverse sequences for a given backbone, reconstructing nucleotide conformations from backbone and preserving functional sites. In benchmarks using experimentally determined ssDNA structures, InvDNA demonstrates more than a twofold improvement in sequence recovery compared with existing ssDNA and RNA sequence design approaches. Further computational validation using AlphaFold3 shows that 44.4% of InvDNA-designed sequences successfully fold into their predefined conformations. Notably, this success rate increases when backbone coordinates are perturbed to diversify the InvDNA-designed sequences. Collectively, these results establish InvDNA as a robust framework for rational ssDNA engineering.

## Introduction

Single-stranded DNA (ssDNA) is an essential biomolecular class involved in key biological processes, including gene regulation^1^, transcriptional control^2^, and the catalysis of biochemical reactions^3,4^. As such, rationally engineered ssDNA can be used to detect and precisely modulate these processes, enabling wide-ranging applications in therapeutics^5,6^, diagnostics^7^, and biosensing^8,9^. Central to these capabilities is the ability to design sequences that reliably adopt a specified backbone conformation, providing a foundation for rational ssDNA engineering^10,11^.

ViennaRNA^12,13^ and NUPACK^14,15^ are the most widely used computational frameworks currently available for ssDNA sequence design. Both approaches approximate the target backbone using secondary structure representations and employ empirical energy functions to identify sequences that achieve a minimum free energy state corresponding to the target secondary structure. However, using secondary structure as the primary design criterion, combined with the reliance on simplified energy functions, presents significant limitations. Secondary structures represent a simplified approximation of backbone geometry, so even if a designed sequence perfectly matches the target secondary structure, its three-dimensional structural fidelity cannot be guaranteed^16,17^. Furthermore, current energy functions approximate complex molecular interactions as linear combinations of a few empirical factors, such as the free energy of specific loop types, and derive their parameters from limited experimental data^18–20^. This oversimplification fundamentally restricts both the accuracy and practical applicability of energy function-based approaches.

In protein and RNA sequence design, which are highly analogous to ssDNA sequence design, deep learning-based methods have largely replaced traditional energy function-based models, establishing a new design paradigm^21–25^. These deep learning-based methods encode backbone structures using detailed geometric descriptors, such as interatomic distances and torsion angles, and employ neural networks to identify sequences that fold into conformations consistent with these encoded geometric features. During training, the neural networks automatically capture high-dimensional correlations between geometric features and sequences. Although these strategies provide valuable guidance for developing ssDNA sequence design methods, their direct application is hindered by the limited availability of ssDNA structural data^26^, relative to that of protein and RNA^22,27^. This data limitation poses a significant challenge to the generalization and robustness of deep learning model^28,29^.

Flexible backbone representations and end-to-end deep learning offer a promising strategy for improving the generalization ability of deep learning-based sequence design models. Unlike fixed geometric features that encode backbones, which restrict the model to a single view of each structure-sequence pair, flexible backbone representations that randomly sample subsets of backbone atoms as encoded backbone enable analysis of each structure-sequence pair from a variety of perspectives. This flexibility has the potential to improve the expressiveness of learned representations and enhance the predictive performance of the model. An end-to-end framework that operates directly on atomic coordinates naturally supports such flexible backbone representation and can also use the full backbone as input, thereby avoiding the loss of structural information during conversion from backbone to geometric features.

The goal of reconstructing all-atom structures from target backbones and designed sequences provides another promising strategy for improving the generalization ability of sequence design models by imposing explicit spatial constraints between designed sequences and their target backbones. Biomolecules with well-defined three-dimensional structures require sequence design methods capable of generating sequences whose all-atom conformations align with the target backbone while satisfying relevant physicochemical constraints. However, current deep learning approaches primarily focus on inferring amino acid or nucleotide identities from geometric features and do not explicitly enforce structural compatibility between designed sequences and target backbones. Several studies, including CarbonDesign^30^, ABACUS-R^23^, and LigandMPNN^31^, incorporate side-chain conformation prediction as an auxiliary training objective and report measurable improvements. Despite these advances, such indirect modelling of spatial relationships does not fully capture local atomic interactions, such as those between backbone and nucleotides atoms, and therefore cannot completely prevent spatial clashes.

In this study, we introduce InvDNA, a deep learning-based method for ssDNA sequence design built on an end-to-end deep learning framework that inversely transforms a predefined backbone conformation into an ssDNA sequence. Specifically, this framework takes backbone atoms and masked sequences as input and outputs designed ssDNA sequences along with all-atom structures. In addition to adopting training strategies including flexible backbone representation and all-atom structure reconstruction objectives, InvDNA incorporates dynamic sequence masking, where 0–20% of nucleotides in the input sequence are randomly retained during training. This retention simulates preserving functionally important nucleotides that may be constrained by their functional roles.

Benchmarking on experimental ssDNA structures shows that InvDNA outperforms traditional energy function-based ssDNA sequence design methods and deep learning-based RNA sequence design approaches, achieving a twofold improvement in sequence recovery rate and a twofold to threefold in the proportion of designed sequences that fold into the target backbone conformation. These results demonstrate the effectiveness of the deep learning paradigm for ssDNA design. Moreover, the training strategies employed by InvDNA not only enhance sequence design accuracy but also enable additional capabilities, including generating diverse sequences for a given backbone, reconstructing nucleotides conformations from backbone and designing sequences that constrain specific nucleotide types at designated positions. Finally, given the limited availability of ssDNA structural data, we investigate how the number of ssDNA structures available for training influences the performance of InvDNA. Our study shows that the performance of InvDNA can be further improved by leveraging additional training data.

## Results

### Overview of InvDNA

The overall architecture of InvDNA is illustrated in Figure 1a. The framework takes masked sequences and backbone coordinates as direct inputs and uses a sequence representation that is iteratively updated through twelve structure blocks to predict both all-atom coordinates and nucleotide identities. For the sequence input, since each position in the ssDNA backbone contains multiple atom types, we model the sequence representation as a three-dimensional tensor that encodes atom-level information across inter-nucleotide, intra-nucleotide, and channel dimensions. The backbone coordinates are also represented using a three-dimensional tensor spanning inter-nucleotide, intra-nucleotide, and spatial dimensions. Coordinates of non-existent atoms (e.g., the “C7” atom in adenine) or atoms that are masked are replaced with the centroid coords of the existing atoms. By default, the backbone uses all backbone atoms for representation (see Figure 1b). When using flexible backbone representation, the atom types “P”, “C3’” and “C1’” are always retained as core atoms, while the remaining atoms are randomly masked to enhance structural diversity during training.

**Figure 1:**
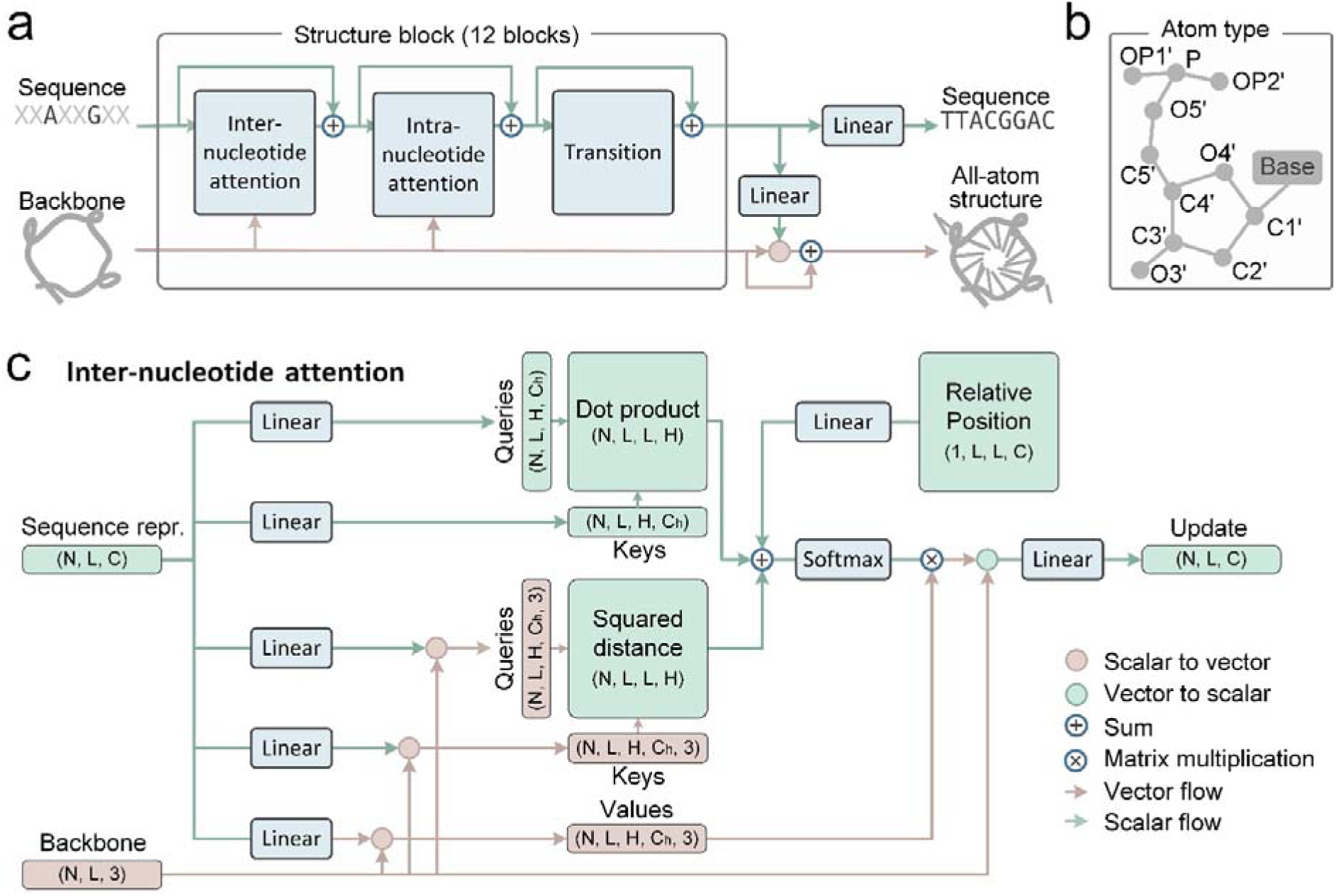
Overview of the InvDNA method. **a**, The end-to-end InvDNA framework comprises 12 structure blocks, each block includes an inter-nucleotide attention module (Figure 1c), an intra-nucleotide attention module (Figure S1), and a transition module^33^. Linear denotes a linear neural network layer. The framework accepts backbone coordinates and, optionally, specified core nucleotides as input. The output is a designed ssDNA sequence alongside its corresponding all-atom structural model. **b**, Visualization of the atom types used for the backbone representation in InvDNA. **c**, Architecture of the inter-nucleotide attention module. The module takes sequence representation and backbone coordinates as input and updates the sequence representation via learned inter-nucleotide interactions. Here, N denotes the number of atoms, L the sequence length, C the number of channels (default 384), and H the number of attention heads (default 12).

Each structure block receives both the sequence representation and backbone coordinates as inputs, while the initial sequence representation is derived from embeddings of the masked sequence. The block then updates the sequence representation using inter-nucleotide attention, intra-nucleotide attention, and transition modules sequentially. As shown in Figure 1c, the inter-nucleotide attention module, inspired by the invariant point attention module proposed by AlphaFold2^32^, updates the sequence representation by considering both the backbone coordinates and the relative nucleotide position within the inter-nucleotide dimension. The intra-nucleotide attention module has a similar architecture to inter-nucleotide attention module (see Figure S1) and updates the sequence representation based on intra-nucleotide interactions within the intra-nucleotide dimension. Finally, the transition module^33^ updates the sequence representation through channel-wise transformations.

### Performance of InvDNA on experimental ssDNA structures

We leveraged a set of recently released experimental ssDNA structures (referred to as the “recent ssDNA dataset”) to benchmark InvDNA against currently available ssDNA sequence design methods, specifically ViennaRNA (using DNA2004 free energy parameters) and NUPACK (using dna04.2 free energy parameters). Given the scarcity of dedicated ssDNA design methods and the structural parallels between nucleic acids, we also evaluated R3Design^21^ and RiboDiffusion^34^, the start-of-the-art deep learning models for RNA sequence design. These models were adapted for ssDNA by substituting uracil (U) with thymine (T) in the output sequences. The recent ssDNA dataset includes 45 ssDNA structures deposited in the Protein Data Bank (PDB)^35^ between December 2021 and June 2024, each sharing less than 100% sequence identity with InvDNA’s training set and exhibiting a length distribution similar to that of the training set. (see Methods and Figure S2). Build on this, we curated an additional non-redundant dataset of 24 ssDNA structures with less than 80% sequence identity to InvDNA’s training set. Given the potential structural-redundancy between the recent ssDNA dataset and the training set, we also prepared another non-redundant dataset of 20 ssDNA structures from the recent ssDNA dataset, each having a TM-score below 0.45 relative to any structure in the InvDNA’s training set. For ViennaRNA and NUPACK, which require secondary structure inputs, we utilized DSSR^36^ to extract secondary structures from the corresponding ssDNA three-dimensional coordinates.

We first evaluated the sequence recovery rates of designs generated by the five methods on both the recent and non-redundant ssDNA datasets. Sequence recovery is defined as the percentage of positions in a designed sequence that match the native sequence for a given backbone^22,37^; higher sequence recovery generally correlates with improved structural fidelity to the target backbone^38^. On both datasets, InvDNA demonstrated superior performance compared to all other methods (Figure 2a, Figure S3-S4). For 76.6% of the targets in the recent ssDNA dataset, the sequences designed by InvDNA exhibited the highest recovery rates, highlighting the efficacy of InvDNA’s deep learning-based approach. Moreover, The performance gap between InvDNA and the deep learning-based RNA sequence design models (R3Design and RiboDiffusion) suggests fundamental differences in the folding landscapes and physicochemical properties of ssDNA versus RNA. Consequently, methods optimized for RNA are not directly transferable to ssDNA, underscoring the necessity of developing specialized tools like InvDNA. Figure 2b and Figure 2c also indicates that InvDNA achieves consistent performance on backbones sharing 80-100% sequence identity with InvDNA’s training set, as those with less than 80% identity, demonstrating the robustness of InvDNA to backbone divergence. Although InvDNA’s performance on the structural-nonredundant test set was slightly lower than on the recent dataset, it remained significantly higher than that of other methods, further confirming its robustness (Figure 2d).

**Figure 2:**
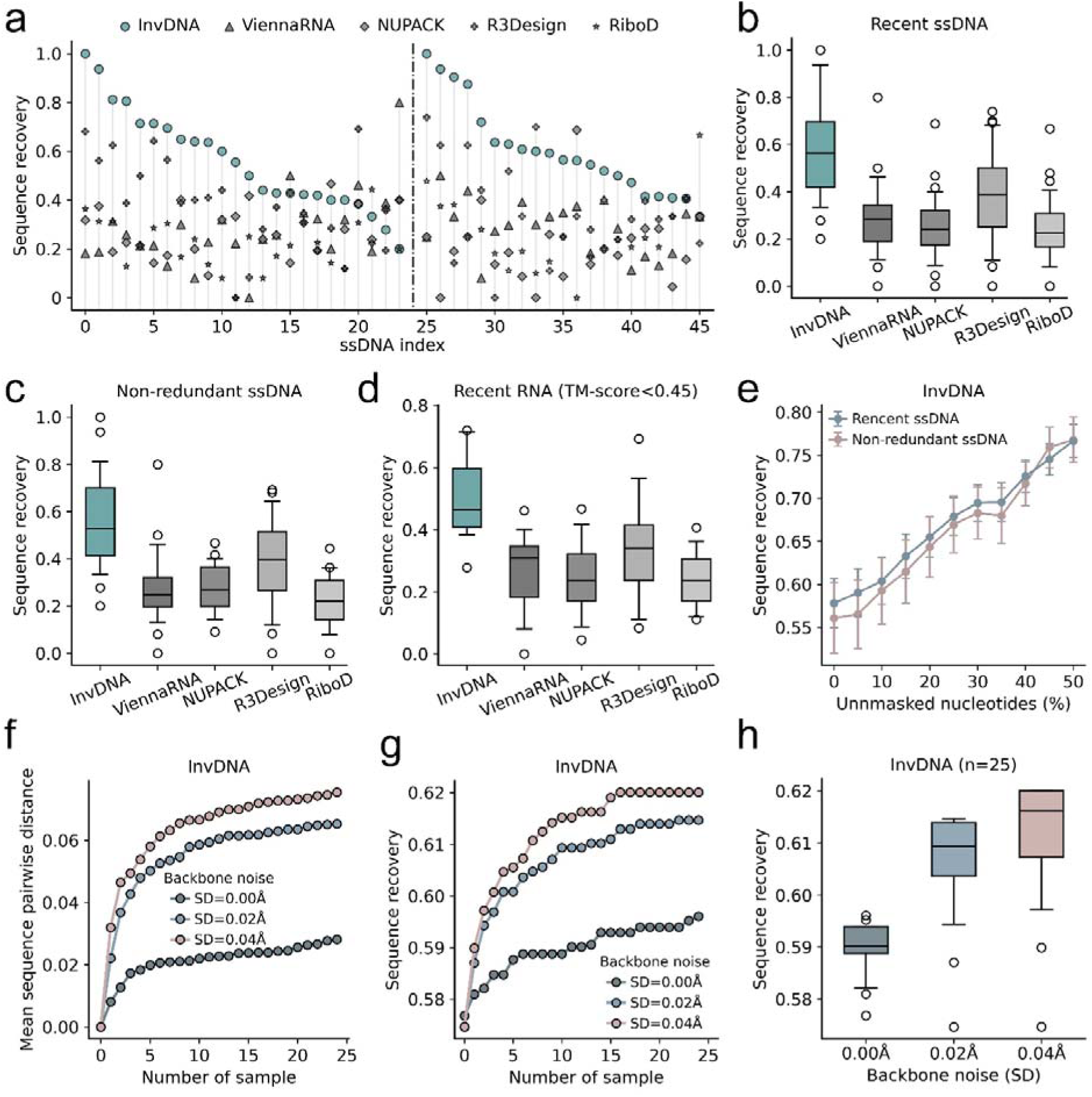
Performance of InvDNA on the ssDNA test set. **a**, Sequence recovery rates for InvDNA, ViennaRNA, NUPACK, R3Design, and RiboDiffusion (RiboD) across each ssDNA backbone in the recent ssDNA dataset. The dashed vertical line separates targets sharing 80–100% sequence identity with the InvDNA training set (left) from those exhibiting <80% identity (right). **b**, Distribution of sequence recovery rates for sequences designed by InvDNA, ViennaRNA, NUPACK, R3Design, and RiboDiffusion (RiboD) on the recent and non-redundant ssDNA datasets. Box limits indicate the 25th–75th percentiles, and the center line represents the median. **c-d**, Distribution of sequence recovery rates for sequences designed by InvDNA, ViennaRNA, NUPACK, R3Design, and RiboDiffusion (RiboD) on the non-redundant ssDNA datasets (**c:** sequence identity of each ssDNA to training set<80%, **d:** tm-score of each ssDNA to training set<0.45). Box limits indicate the 25th–75th percentiles, and the center line represents the median. **e**, Sequence recovery rates for sequences generated by InvDNA as a function of the proportion of randomly retained nucleotides in the input sequences. The error bars represent standard deviation, standard error. **f**, Pairwise distance of sequences generated by InvDNA as a function of the total number of sequences generated. Sequences were generated using a flexible backbone representation perturbed by varying levels of Gaussian noise (standard deviation = 0.00 Å, 0.02 Å, 0.04 Å). **g**, Sequence recovery rates of InvDNA as the number of generated sequences increases. For each ssDNA backbone, the sequence with best recovery in the total generated sequences was selected as representative. **h**, Distribution of sequence recovery rates for sequences designed by InvDNA on the recent ssDNA dataset using flexible backbone representation with different Gaussian noise (standard deviation=0.00 Å, 0.02 Å, 0.04 Å). For each ssDNA backbone, 25 sequences were generated, and the sequence with the highest recovery was selected as representative.

To investigate the ability of InvDNA to preserve specified nucleotides as one of its training objectives, we analyzed the sequence recovery rate of sequences designed by InvDNA while systematically varying the proportion of unmasked nucleotides in input sequence (from 0% to 50%). As shown in Figure 2d, the sequence recovery rate of InvDNA increases nearly linearly with the percentage of unmasked nucleotides, reaching 99.2% when the complete native sequence is provided as input, demonstrating that InvDNA can effectively incorporate partial sequence constraints.

Given the importance of generating diverse sequences for a single backbone, we further assessed InvDNA’s capacity to produce sequence diversity. InvDNA was trained with flexible backbone representations that perturb the atom-type encoding of the input backbone, thereby naturally enabling variation in the resulting sequences. Additionally, introducing Gaussian noise into the backbone atom coordinates offers an alternative perturbation strategy, an approach previously used in ProteinMPNN^22^ to enhance design diversity. For each backbone in the recent ssDNA dataset, we generated 25 sequences using the flexible-backbone strategy with varying levels of Gaussian noise. As shown in Figure 2f-h, both the mean pairwise distance of designed sequences and mean sequence recovery increases steadily as the number of sampled sequences grows (the mean number of unique sequences is shown in Figure S5, defined as those sharing less than 100% pairwise identity). Notably, perturbing the backbone coordinates with Gaussian noise (standard deviation = 0.02 Å) substantially improves both sequence recovery rate and sequence diversity compared to using flexible-backbone representations alone (standard deviation = 0.0 Å). We also evaluated a larger noise magnitude (standard deviation = 0.04 Å); however, this yield only marginal improvements ( 1% in sequence recovery, also see Figure S6). The distribution of sequence recovery rates for each configuration on the recent ssDNA dataset is shown in Figure 2g, demonstrating that sequence recovery improves with increasing noise levels before eventually plateauing. Together, these results demonstrate that InvDNA effectively produces multiple diverse designs for a given backbone, providing a rich pool of candidates for downstream experimental screening and optimization.

### In silico evaluation of InvDNA using AlphaFold3

We further assessed whether the sequences designed by InvDNA could fold into their intended backbone conformations. The three-dimensional structures of the designed ssDNA sequences were predicted using AlphaFold3^33^, a leading structure prediction model for biomolecular complexes. As shown in Figure 3a, InvDNA generate one designed sequence per backbone, the structural similarity between each AlphaFold3-predicted structure and its corresponding target backbone was quantified using three metrics: the Local Distance Difference Test (LDDT), the Global Difference Test – Total Score (GDT-TS), and the root-mean-square deviation of C3’ atoms (RMSD-C3’). LDDT evaluates local structural accuracy by assessing the consistency of interatomic distances within a defined neighborhood; an LDDT score of 1.0 indicates perfect agreement between the predicted and reference structures. GDT-TS measures the percentage of nucleotide positions falling within a defined distance threshold of the reference structure, with higher values reflecting greater global structural similarity. C3’-RMSD quantifies the average displacement of C3’ atoms between the predicted and reference structures, where lower values indicate closer structural agreement. It is worth noting that All ssDNA in the recent dataset were released after March 2022 (see Table S1), ensuring no overlap with the AlphaFold3 training set. In addition to the standard InvDNA output, we benchmarked the structures predicted from the sequences designed by ViennaRNA, NUPACK, R3Design, RiboDiffusion, and InvDNA with flexible backbone representations and Gaussian noise (standard deviation = 0.04 Å).

**Figure 3:**
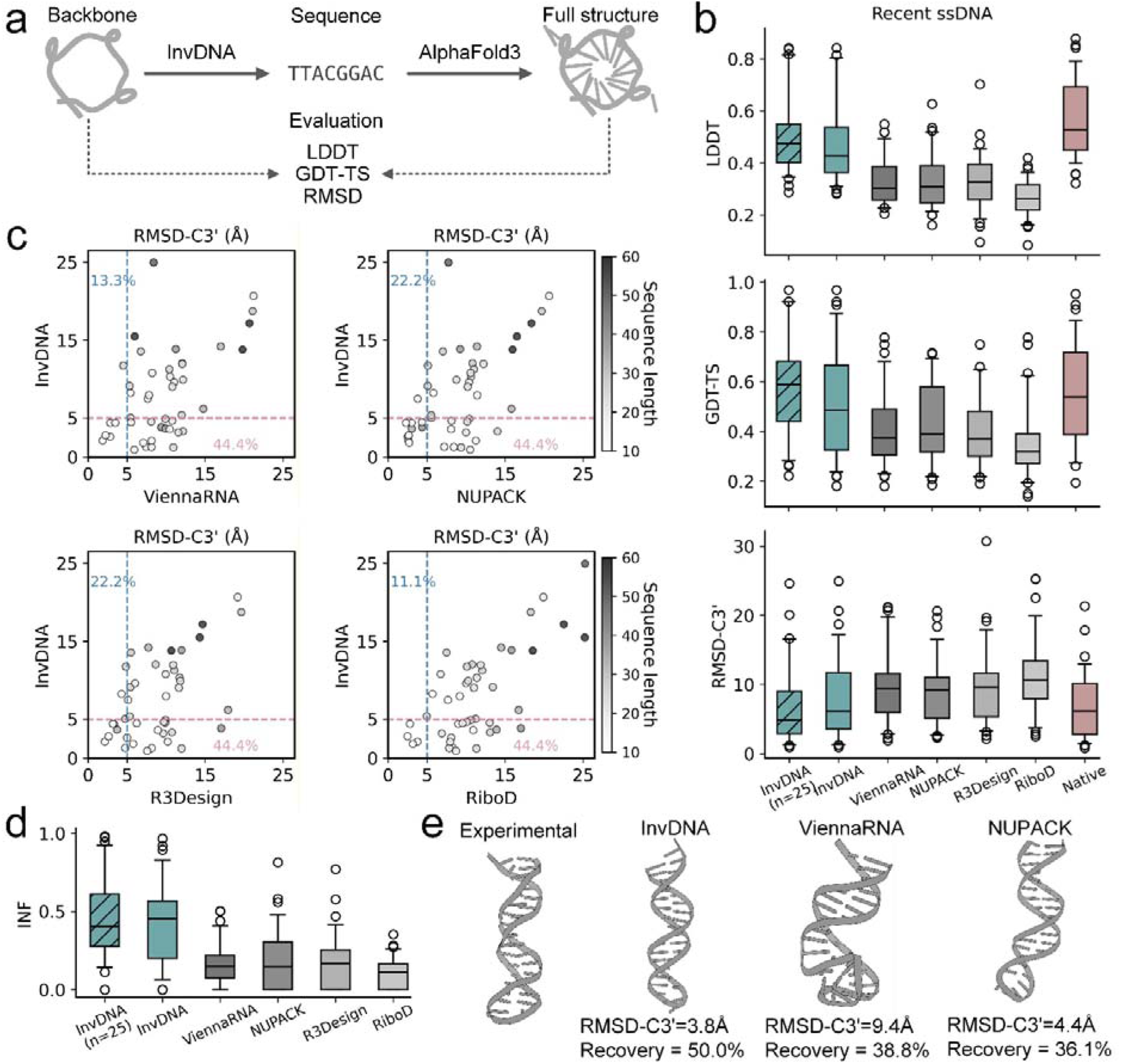
In silico evaluation of InvDNA using AlphaFold3. **a**, Workflow for evaluating the structural consistency between structures folded from designed sequences and the intended target backbone. **b**, Distribution of LDDT, GDT-TS, and RMSD-C3’ scores for structures predicted by AlphaFold3. Predictions were performed on the recent ssDNA dataset using native sequences and sequences generated by InvDNA, ViennaRNA, NUPACK, R3Design, and RiboDiffusion. For InvDNA (n=25), 25 sequences were generated per ssDNA backbone using a flexible backbone representation with Gaussian noise (standard deviation = 0.04 Å); the sequence yielding the predicted structure with the highest confidence was selected as representative. **c**, Pairwise comparisons of RMSD-C3’ scores between structures predicted from InvDNA-designed sequences (y-axis) and those from ViennaRNA, NUPACK, R3Design, or RiboDiffusion (x-axis). All structures were predicted using AlphaFold3. **d**, Distribution of INF scores for structures predicted by AlphaFold3. Predictions were performed on the recent ssDNA dataset using sequences generated by InvDNA, ViennaRNA, NUPACK, R3Design, and RiboDiffusion. For InvDNA (n=25), 25 sequences were generated per ssDNA backbone using a flexible backbone representation with Gaussian noise (standard deviation = 0.04 Å); the sequence yielding the predicted structure with the highest confidence was selected as representative. **e**, Representative structures predicted by AlphaFold3 using sequences designed by InvDNA, ViennaRNA, NUPACK, R3Design, and RiboDiffusion for a hairpin backbone (PDB: 8BM6). All predicted structures are displayed aligned to the experimental backbone.

Figure 3b displays the distributions of LDDT, GDT-TS, and RMSD-C3’ values for each method on the recent ssDNA dataset. Across all three metrics, InvDNA significantly outperformed ViennaRNA, NUPACK, R3Design, and RiboDiffusion. Notably, incorporating flexible backbone representations and Gaussian noise into InvDNA to generate diverse sequences further enhanced the similarity between AlphaFold3-predicted structures and target backbones across all metrics. Given the limitations of AlphaFold3, where even when provided with native sequences, only a subset of ssDNA (18 of 45 ssDNA have RMSD-C3’ less than 5 Å) successfully adopt their native conformations, we re-analyzed the performance of each sequence design methods on these 18 ssDNA (see Figure S7). The quality of the predictions generated by AlphaFold3 from InvDNA-generated sequences remains significantly superior to that of other methods. To quantify folding success, we defined a successful folding design as one achieving an RMSD-C3’ value below 5 Å (see Table S2 for detailed values). As shown in Figure 3c, the success rate of sequences designed by InvDNA (without sampling) reached 44.4%, approximately fourfold higher than that of RiboDiffusion (11.1%), threefold higher than that of ViennaRNA (13.3%) and twofold higher than NUPACK and R3Design (each 22.2%).

Since ViennaRNA and NUPACK was originally designed for generated sequences that can refold into the desired secondary structure, we also evaluated the quality of the predicted structures using the Interaction Network Fidelity (INF) metric. INF was calculated using RNAassessment^39^ to quantitatively evaluate the structural accuracy of the predicted structure by measuring the recapitulation of native canonical and non-canonical interaction networks. An INF value of 1.0 indicates perfect agreement with the reference structure, while a value of 0 suggests complete mismatch. As shown in Figure 3d, structures predicted by AlphaFold3 using InvDNA-generated sequences exhibited significantly higher INF values than those from ViennaRNA and NUPACK, indicating that sequences designed by InvDNA also have an advantage in recreating the secondary structure. A representative example is the sequence designed for the hairpin (PDB: 8BM6). As shown in Figure 3d (see Figure S8 for more examples), the structure predicted from the InvDNA-designed sequence attained an RMSD-C3’ of 3.8 Å, whereas sequences designed by ViennaRNA, NUPACK yielded RMSD-C3’ values of 9.4 Å, 4.4 Å, respectively. Together, these results demonstrate that InvDNA achieves substantially higher structural fidelity than existing methods and highlight the utility of combining InvDNA with AlphaFold3 as a computational pipeline for ssDNA sequence design.

### Reconstruction of all-atom structure from backbone

Accurate reconstruction of nucleobase conformations (analogous to side-chain packing in proteins) is a crucial step in molecular structure prediction^40–42^. Although the primary InvDNA model was not explicitly designed to reconstruct nucleobase conformations from backbones and native sequence information, the convergence of the structural reconstruction loss during training and its ability to retain the identities of key nucleotides suggest that it may inherently be capable of reconstructing full atomic coordinates. We therefore tested the ability of InvDNA to reconstruct nucleobase conformations when both the complete native sequence and backbone are provided, evaluating performance on the recent ssDNA dataset (Figure 4a). Alongside results from the knowledge-based method PDBFixer^43^ and the deep learning-based method StruCloze^44^ as a control, we trained a dedicated InvDNA version (denoted as InvDNA (full sequence)) to reconstruct nucleobase conformations by taking the complete sequence and backbone coordinates as input during training.

**Figure 4:**
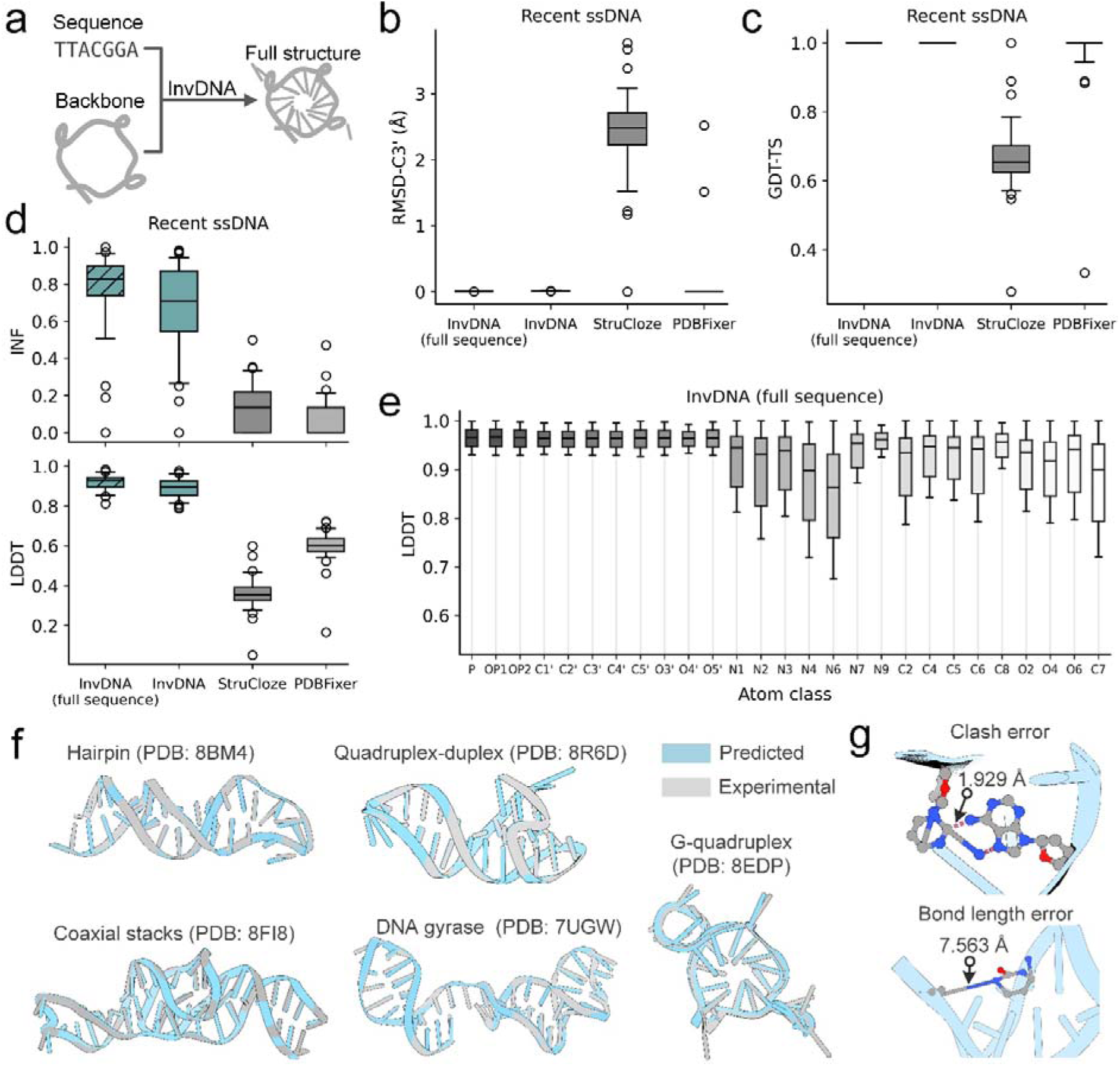
Reconstruction of all-atom structures from backbone coordinates. **a**, Workflow for evaluating structural consistency between InvDNA-predicted all-atom structures and experimental references. **b-c**, Distributions of RMSD-C3’ (**b**) and GDT-TS (**c**) scores for structures predicted by InvDNA (full sequence), InvDNA, StruCloze, and PDBFixer. Predictions were performed on the recent ssDNA dataset using full sequences and full backbones as input. InvDNA (full sequence) denotes a re-trained version of InvDNA that use the full sequence as input and a structure-loss-only training objective. **d**, Distribution of INF and LDDT scores for structures predicted by InvDNA (full sequence), InvDNA, StruCloze, and PDBFixer. **e**, Per-atom LDDT values comparisons between experimental structures and those generated by InvDNA (full sequence). **f**, Example of InvDNA (full sequence)-generated all-atom structures (cyan) and the corresponding experimental structures (gray). Nucleotides are depicted as stubs. **g**, Local structural details of atomic clashes and bond-length deviations in structures predicted by InvDNA (full sequence).

We first evaluated the backbone reconstruction accuracy of the four methods relative to the target experimental structures using RMSD-C3’ and GDT-TS. As shown in Figure 4b-c, the backbone of the all-atom structures generated by InvDNA (full sequence), the original InvDNA, and PDBFixer were nearly identical to the target backbones. In contrast, structures generated by StruCloze exhibited a backbone error of approximately 2 Å.

Next, we assessed the fidelity of the reconstructed nucleobase conformations using Interaction Network Fidelity (INF) and LDDT. As shown in Figure 4d, the nucleobase conformation accuracy of all-atom structures generated by InvDNA and InvDNA (full sequence) was significantly higher than that of both StruCloze and PDBFixer, demonstrating the practicality of InvDNA. As expected, the InvDNA (full sequence) model showed systematic improvements over the original InvDNA. It is worth noting that the median INF of InvDNA (full sequence) reached 0.8, indicating that the predicted all-atom structures possess reasonable hydrogen bond networks. The per-atom quality of the all-atom structures generated by InvDNA (full sequence) is detailed in Figure 4e. The median LDDT for atoms within the nucleobases reached nearly 0.9, indicating that most nucleobase atoms were placed at correct positions, while the median LDDT for backbone atoms exceeded 0.95, reflecting extremely high fidelity to the experimental backbone.

Figure 4f illustrates structures generated by InvDNA (full sequence) using the complete backbone and native sequence as inputs. Across ssDNA targets with varying structural topologies and functions, the nucleobases in most predicted structures adopted the correct orientation relative to the native structure, further confirming the positional accuracy of the atoms. It should be noted that the structures generated by InvDNA (full sequence) may contain minor physical inconsistencies, such as steric clashes or bond length deviations (see Figure 4g). However, these artifacts can be effectively resolved by coupling the generation process with molecular dynamics-based refinement^43^.

### Interpreting the InvDNA

We trained three ablation models to evaluate the contribution of three key components involved in training InvDNA: flexible backbone representations, structure reconstruction objectives, and dynamic nucleotide masking. Specifically, the “No atom sampling” model was trained by replacing the flexible backbone representation with full backbone atoms as input. The “No structure loss” model was trained by removing structure reconstruction related losses, including clash loss, bond loss and all-atom frame aligned point error (FAPE) loss (see Methods). The “No dynamic masking” model was trained by removing the strategy of random retaining 0-20% of nucleotides in input sequence during training, resulting in a fully masked sequence throughout training. After convergence, all models were evaluated on the recent ssDNA dataset using fully masked sequence and complete backbone atoms as input during inference.

We first investigated the sequence recovery rate and RMSD-C3’ of AlphaFold3-predicted structures for sequences generated by InvDNA and the three ablation models, as shown in Figure 5a (see Figure S9 for pairwise comparisons). As hypothesized, removing flexible backbone representations or structure reconstruction objectives consistently decreased both sequence recovery rate and RMSD-C3’, demonstrating that each component improves the generalization capability of sequence design model. Notably, the dynamic nucleotide masking strategy, which aims to simulates preserving functionally important nucleotides, also slightly enhances the model’s generalization ability. Specifically, InvDNA achieves median sequence recovery and RMSD-C3’ of 56.3% and 6.18 Å, respectively, compared to 51.9% and 8.29 Å for "No atom sampling," 50.0% and 7.92 Å for "No structure loss," and 54.5% and 8.90 Å for "No dynamic masking." Mean values for each model are reported in Table S2.

**Figure 5:**
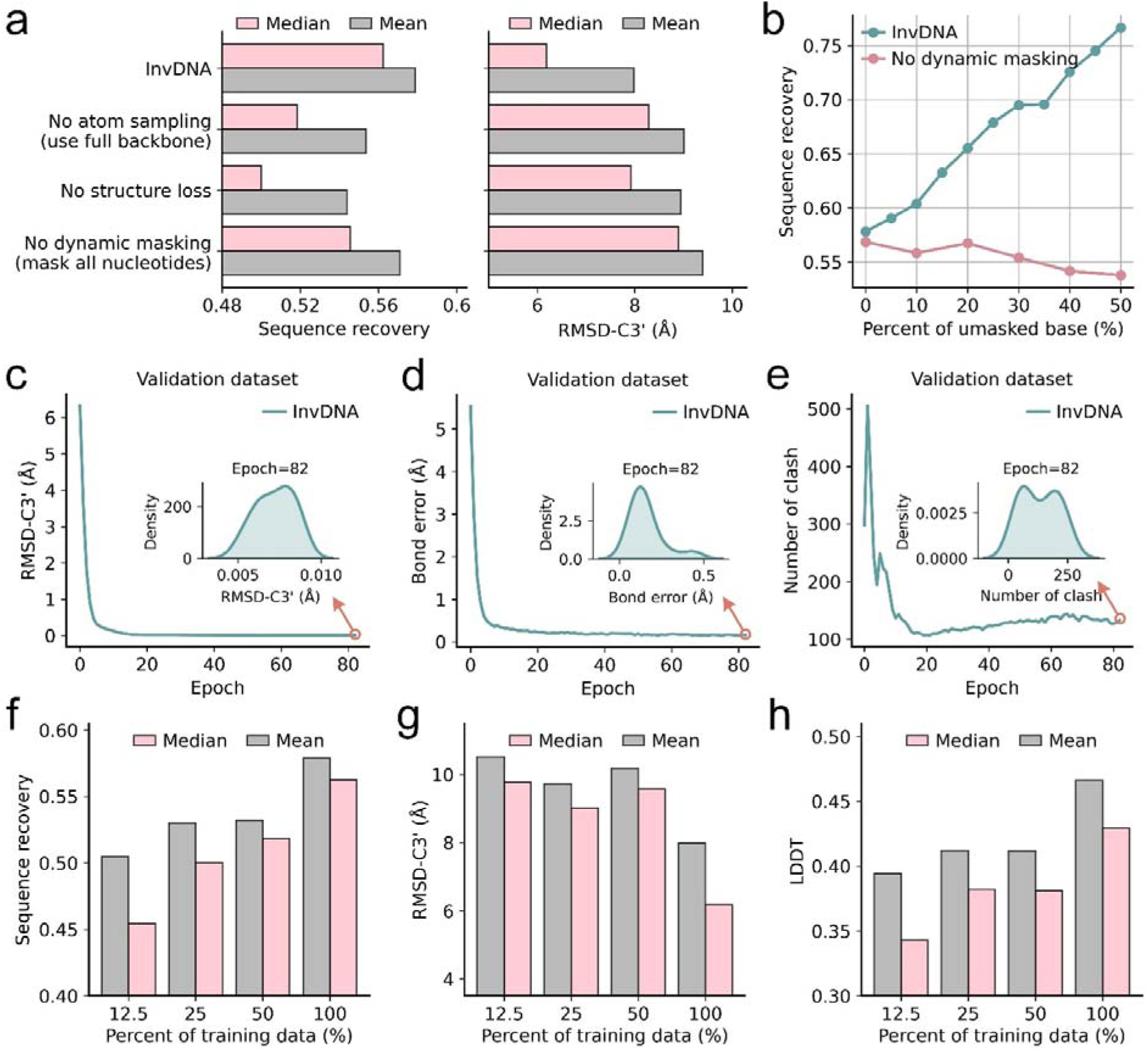
Ablation study of InvDNA training strategies. **a**, Mean and median sequence recovery rates for InvDNA and ablated variants on the recent ssDNA dataset. Structures predicted using AlphaFold3 from these generated sequences are evaluated using RMSD-C3’. **b**, Sequence recovery rates for InvDNA and the “No dynamic masking” model as a function of the proportion of randomly retained nucleotides. **c-e**, Training curves for InvDNA on the validation dataset, quantified by RMSD-C3’ (**c**), bond-length error (**d**), and Number of atomic clashes (**e**). **f**, Mean and median sequence recovery rates for InvDNA and ablated variants that trained with different training data size on the recent ssDNA dataset. **g-h**, Structural evaluation of sequences generated by models trained on varying data sizes. AlphaFold3 predictions are assessed using RMSD-C3’ (**g**) and LDDT (**h**).

To investigate the ability of InvDNA to preserve specified nucleotides as one of its training objectives, we analyzed the sequence recovery rates of sequences designed by InvDNA and the “No dynamic masking” model while systematically varying the proportion of unmasked nucleotides in the input (from 0% to 50%). As shown in Figure 5b, InvDNA’s sequence recovery rate increased nearly linearly with the percentage of unmasked nucleotides, reaching 99.2% when the complete native sequence was provided as input. In contrast, the "No dynamic masking" model, which was trained on fully masked input sequences, was unable to effectively utilize the provided sequence information, resulting in a slight decrease in sequence recovery rate as more nucleotides were unmasked. This result demonstrates that the dynamic nucleotide masking strategy is essential for the model to effectively incorporate partial sequence constraints.

We also investigated the ability of InvDNA to reconstruct physically realistic all-atom structures from the backbone, as another key training objective. The quality of all-atom structures reconstructed during training on the validation dataset is shown in Figure 5c-e (mean sequence cross-entropy loss and sequence recovery rate are also shown in Figure S10). Specifically, Figure 5c shows the mean RMSD-C3’ of reconstructed structures relative to the input backbone, while Figure 5d-e show the mean bond length deviation and number of steric clashes, respectively. All three metrics improve substantially during training: the RMSD-C3’ decreases from an initial value of 6.5 Å to 0.007 Å, bond length deviation decreases from 5.8 Å to 0.14 Å, and the number of steric clashes decreases from nearly 500 to nearly 175 per prediction. This substantial improvement confirms that the structure reconstruction-related losses effectively guide the model in predicting physically plausible all-atom structures from the backbone.

Given the limited availability of ssDNA structural data, we also investigated how the amount of available ssDNA structures for training affects the performance of InvDNA. Specifically, we employed the same training methodology as InvDNA to train three additional ablation models, each using randomly selected 12.5%, 25%, and 50% of the total clusters from the training data. The ssDNAs in the training dataset of InvDNA were clustered using CD-HIT^45^ with sequence identity 80%, resulting into 255 clusters. After convergence, all models were evaluated on the recent ssDNA dataset using fully masked sequence and complete backbone atoms as input during inference.

Figure 5f shows the median and mean sequence recovery rates of sequences generated by InvDNA and the three data-scaling ablation models. Performance improved consistently as the amount of available ssDNA structural data increases, demonstrating that InvDNA’s performance is currently constrained by the quantity of available ssDNA structures. We further evaluated these models by assessing the quality of AlphaFold3-predicted structures using the generated sequences. As shown in Figure 5g-h, the mean LDDT (calculated using C3’ atoms) and mean RMSD-C3’ for the sequence design models also increase with more available ssDNA structural data. Pairwise comparisons between the ablation models and InvDNA (shown in Figure S11) further confirm that these increases were systematic. Median and mean values for sequence recovery rate, LDDT and RMSD-C3’ values for each model are reported in Table S3. Notably, even when trained on just 12.5% of the full training clusters, InvDNA still significantly outperforms traditional methods in sequence recovery rate, underscoring the effectiveness of deep learning approaches for ssDNA sequence design.

## Discussion

In this work, we present InvDNA, a deep learning framework for ssDNA sequence design that takes backbone atoms and masked sequences as input and outputs designed ssDNA sequences alongside all-atom structures. Several training strategies — including flexible backbone representations, structure reconstruction objectives, and dynamic nucleotide masking — are incorporated to enhance sequence generation, support partial sequence constraints, and enable all-atom structure reconstruction from backbone coordinates.

Benchmarking on experimentally determined ssDNA structures demonstrate that InvDNA consistently outperforms existing ssDNA sequence design methods across multiple evaluation scenarios. Ablation experiments confirm that each training component contributes meaningfully to overall performance. Beyond sequence design, InvDNA can generate diverse and structurally viable sequences for a given backbone, accurately reconstruct nucleobase conformations from the backbone, and preserve user-specified nucleotides in the input sequence. We also find that InvDNA’s performance scales with the availability of training data, yet even when trained on a fraction of available structures, it surpasses traditional methods in sequence recovery rate — highlighting the capacity of deep learning to extract meaningful structural patterns from limited ssDNA data.

Reliable operation of InvDNA still depends on a complete backbone, and minor physical inconsistencies remain in the reconstructed all-atom structures. Addressing these challenges presents an opportunity to further improve the generalization and applicability of InvDNA. Additionally, the present study is limited to computational evaluation; wet-lab validation will be essential for a comprehensive assessment of InvDNA’s reliability in practice, and experimental feedback may reveal specific failure modes and opportunities for further improvement that are not apparent from in silico benchmarking alone. Besides, Although InvDNA is primarily designed for ssDNA sequence design, its ssDNA-independent framework and training strategies allow it to be easily extended to protein, RNA, and other biological sequence design tasks.

## Methods

### Training and validation datasets

All DNA chains deposited in the Protein Data Bank (PDB) prior to 30 December 2020 were retrieved to construct the training dataset for InvDNA. Given the scarcity of single DNA chain with stable conformations (i.e., those not in close spatial proximity (within 10.0 Å) to other biomolecules), as most DNA chains in PDB exist in complex form, we collected DNA chains with at least locally stable conformations to construct the training dataset (similar to NuFold^46^). Specifically, DNA chains were retained if they met the following criteria: (1) resolution better than 9 Å if a reported resolution was available; (2) no non-standard residues; (3) at least four resolved residues; and (4) at least four non-local contacts. A non-local contact is defined as minimal interatomic distance (heavy atoms) between nucleotide i and nucleotide j of less than 5 Å, with |j-i| greater than 3. All the collected structures with identical sequences were clustered, yielding 480 unique DNA sequences. This collection, referred to as the "experimental dataset", served as the training dataset for InvDNA.

DNA chains deposited in the PDB between 30 December 2020 and 30 December 2021 were used to construct the validation dataset. After applying the same filtering criteria as for the experimental dataset, the remaining chains were clustered with those from the experimental dataset using CD-HIT-EST-2D (-i2 validation.fasta -i train.fasta -c 1 -M 0 -T 0 -g 1 -n 2 -d 60 -G 1 -r 0 -l 1) at 100% sequence identity to eliminate redundancy. From clusters composed exclusively of candidate validation DNA chains, reference monomers were retained. This process resulted in a validation dataset comprising 27 unique DNA sequences. It is worth noting that, for DNA structures in both the training and validation dataset that without structure in forms of complexes, nucleotides lacking non-local contacts were masked during model training.

### Test dataset

DNA chains deposited in the PDB between 30 December 2021 and 17 June 2024 were used to construct the test dataset. After applying nearly the same filtering criteria as for the experimental dataset, with the exception of a stricter requirement that at least 80% of the base sites in the DNA have at least one non-local contact, the remaining chains were clustered with those from both the experimental and the validation dataset using CD-HIT-EST-2D (-i2 test.fasta -i train+valid.fasta -c 1 -M 0 -T 0 -g 1 -n 2 -d 60 -G 1 - r 0 -l 1) at 100% sequence identity to eliminate redundancy. From clusters composed exclusively of candidate test DNA chains, reference monomers were retained. This process resulted in a test dataset comprising 45 unique DNA sequences. The non-redundant ssDNA dataset

### Training protocol

We trained InvDNA and ablation models on an H100 GPU with automatic mixed precision, exponential moving average, and a batch size of 1. The AdamW optimizer, with a weight decay of 0.01 and a learning rate of 0.001, was used for model optimization. The model was trained for up to 100 epochs and the weights that exhibited the best performance on the validation dataset, based on sequence cross-entropy loss, were saved as the final model. During training, ssDNAs were randomly sampled from the experimental dataset. To improve training stability, we applied gradient clipping by global norm with a threshold of 0.1 on the parameters for each training example.

The loss function for InvDNA consists of sequence-related cross-entropy loss and structure-related losses, including clash loss, bond loss and FAPE loss^32^. The cross-entropy loss aimed to restore native sequences: 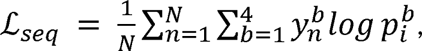 denote one-hot encodings of nucleotide, and the bin probabilities 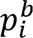 are obtained by linearly project the sequence representations into 4 bins and normalization using a softmax function. The clash loss penalizes inter-atom distances shorter than a defined tolerance: 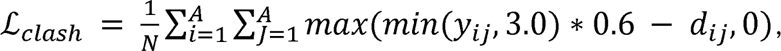 where N is the number of nucleotides in the sequence, A is the number of heavy atoms in the structure. and *y_ij_* and *d_ij_* denote the interatomic distances in the experimental and predicted structure, respectively. The tolerance of atom overlap is set to 0.6. The bond loss enforces bond lengths within each nucleotide to remain within a reasonable length: 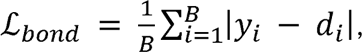 where B is the number of bonds in the structure. and *y_i_* and *d_i_* denote the bond lengths in the experimental and predicted structure, respectively. Considering that the convergence rate of structural losses is much slower than that of sequence loss, we reduced the weight of sequence loss. The corresponding weights for each loss component are set as 0.01 for cross-entropy loss, 0.05 for the clash loss, 0.05 for the bond loss, and 1 for the FAPE loss.

### Evaluation Metrics

The quality of the designed sequence and the AlphaFold3-predicted or InvDNA-predicted structures using designed and native sequence was assessed using the sequence recovery rate, pairwise distance, LDDT, INF, GDT-TS and RMSD metrics.

The sequence recovery rate was calculated according to:

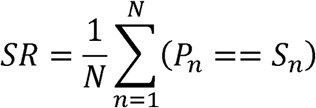

Where N is the number of positions in backbone, P denote the designed sequence and S denote the native sequence. The pairwise distance of designed sequences was calculated according to:

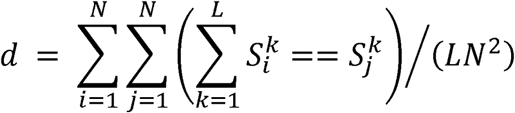

Where N is the number of sequences, L is the number of positions in backbone, 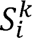 denote the nucleotide of k position in the ith designed sequence. The LDDT, GDT-TS, and RMSD between predicted structures and predefined backbones was evaluated by the OpenStructure^47^ package. The INF was evaluated by the RNAassessment^39^ package.

### Benchmark Methods

ViennaRNA was collected from the latest version, 2.7.0, available on GitHub: https://github.com/ViennaRNA/ViennaRNA. NUPACK was collected from the latest version 4.0.2.0, available at https://www.nupack.org/download/software. AlphaFold3 was collected from GitHub: https://github.com/google-deepmind/alphafold3. RiboDiffusion was collected from GitHub: https://github.com/ml4bio/RiboDiffusion/. R3Design was collected from GitHub: https://github.com/A4Bio/R3Design/tree/master/R3Design. PDBFixer was collected from GitHub: https://github.com/openmm/pdbfixer. StruCloze was collected from GitHub: https://github.com/Junjie-Zhu/StruCloze. All benchmark methods were performed using standard settings.

## Acknowledgements

We thank the developers and contributors of open-source tools and datasets including PDB, PyTorch and others for greatly accelerating our research. This work was supported by National Key R&D Program of China (2022YFA1004800, 2025YFF1207900), Natural Science Foundation of China (T2350003, T2341007, 12131020, 42450084, 42450135, 12326614, 12404236 and 12426310), Zhejiang Province Vanguard Goose-Leading Initiative (2025C01114), Science and Technology Commission of Shanghai Municipality (23JS1401300), and Hangzhou Institute for advanced study of UCAS (2024HIAS-P004), and JST Moonshot R&D (JPMJMS2021).

## Author contributions

L.C. and Y.S. conceived the study. Y. S. performed the experiments and analyzed the results. L.C. and Y.S. wrote the manuscript. Y.X. developed the web server for InvDNA. L.C. supervised the research and provided funding.

## Data and Code availability

The source code and pretrained weights of InvDNA have been released at https://github.com/yunda-si/InvDNA and https://biostructure.sibcb.ac.cn.

## Supplementary Information

### Figures

**Figure S1:**
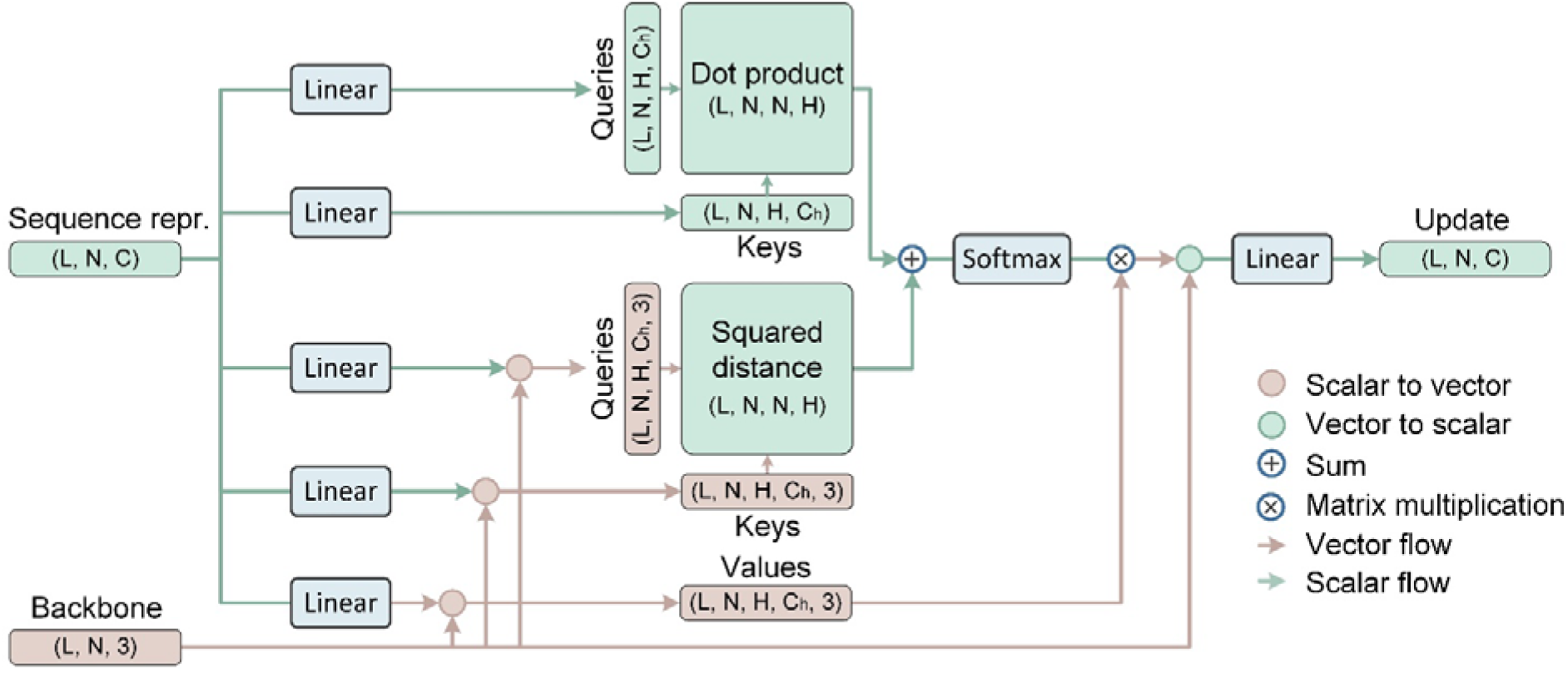
Architecture of the intra-nucleotide attention module. Linear refers to linear layer in neural network. The module takes sequence representation and backbone coordinates as input and outputs an updated sequence representation by learning intra-nucleotide interactions. Here, N denotes the number of atoms, L the sequence length, C the number of channels (default 384), and H the number of attention heads (default 12).

**Figure S2:**
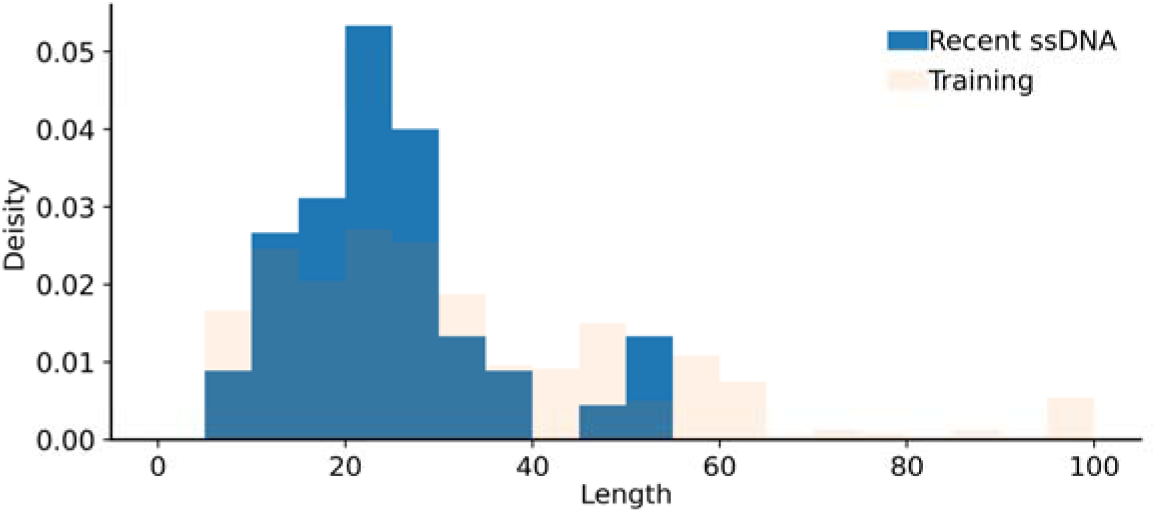
Length distribution of ssDNA in training dataset and recent ssDNA dataset.

**Figure S3:**
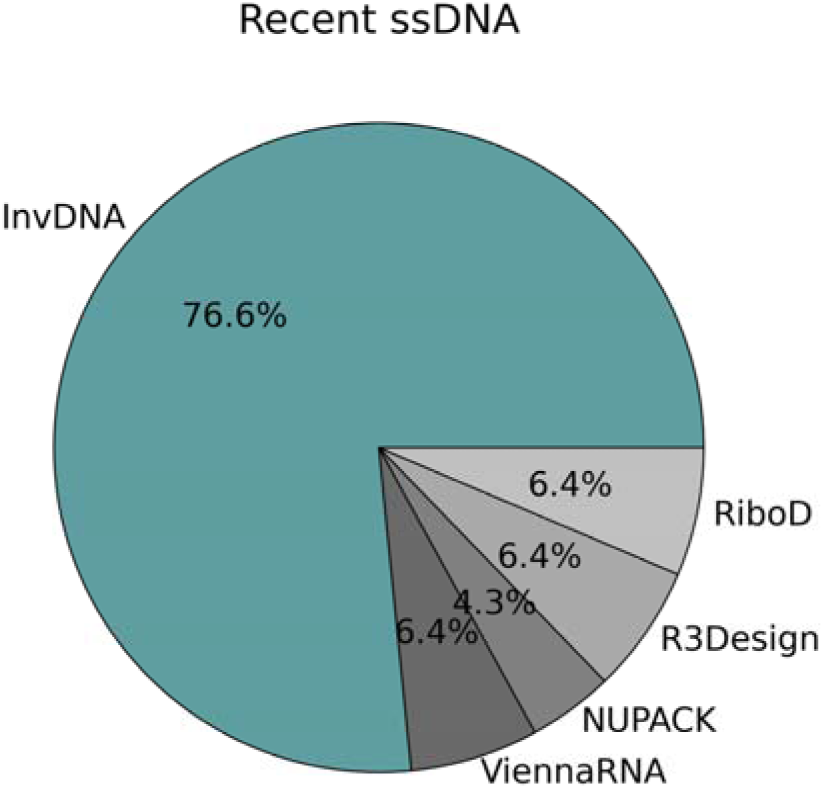
Percentage of ssDNAs for which each method achieves the highest sequence recovery rate.

**Figure S4:**
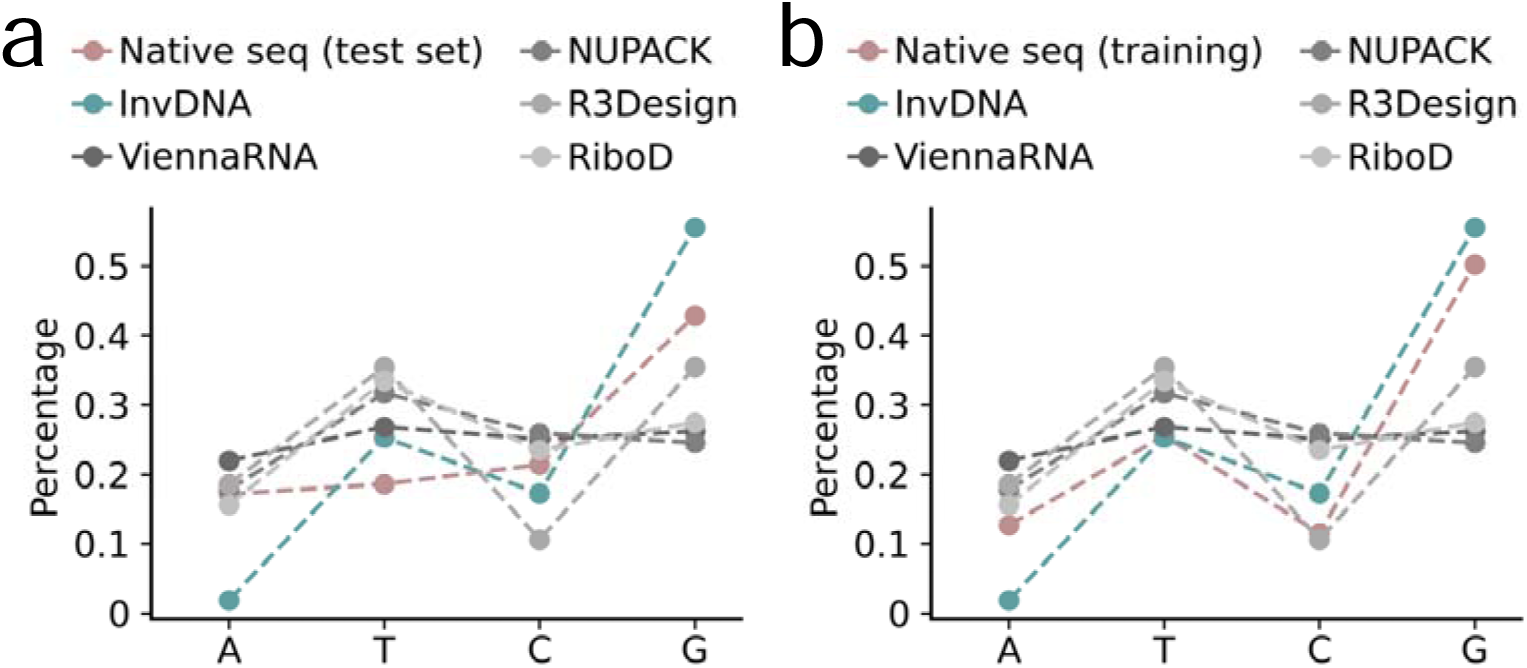
Distribution of nucleotide types of sequences designed by InvDNA, NUPACK, ViennaRNA, R3Design, RiboD on the recent ssDNA dataset.

**Figure S5:**
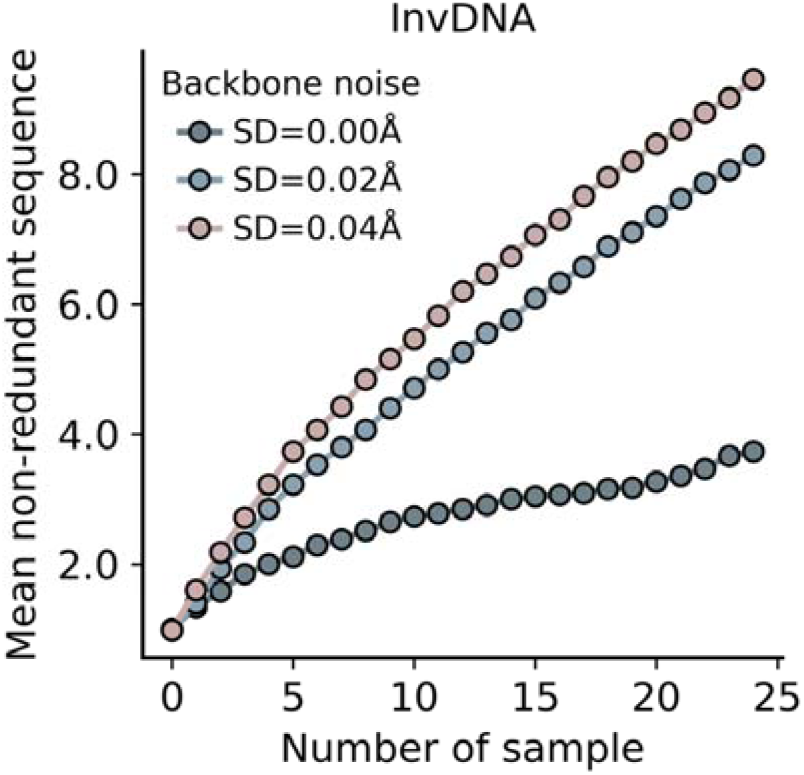
The number of non-redundant sequences (pairwise sequence identity <100%) generated by InvDNA as a function of the total number of sequences generated. Sequences were generated using a flexible backbone representation perturbed by varying levels of Gaussian noise (standard deviation = 0.00 Å, 0.02 Å, 0.04 Å).

**Figure S6:**
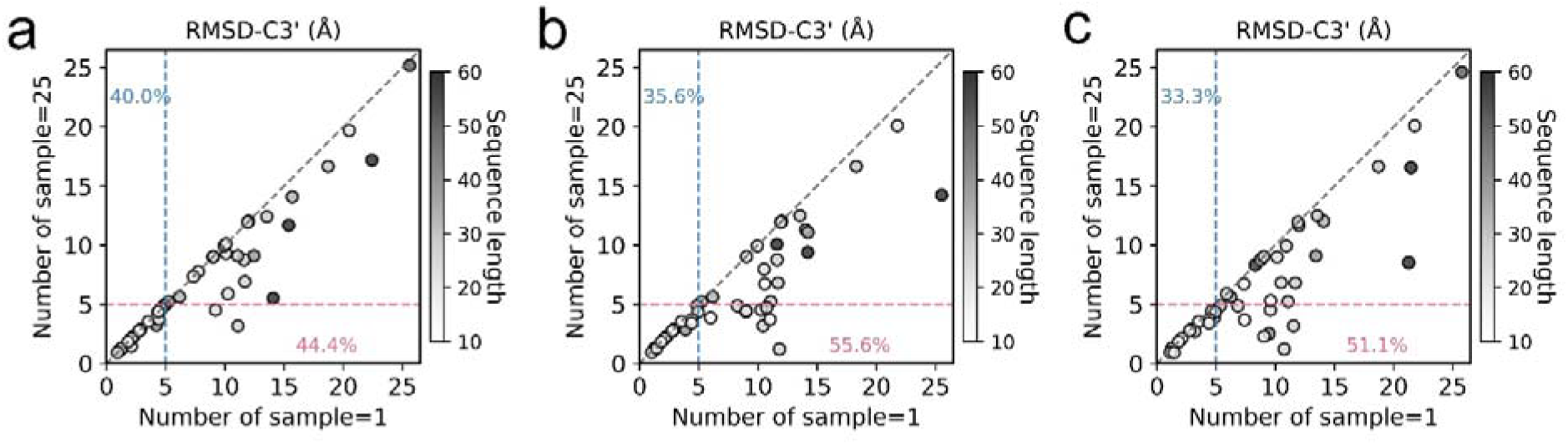
Pairwise comparisons between the best-performing predicted structures using InvDNA-generated sequences (x-axis: 1 generated sequence per backbone; y-axis: 25 generated sequences per backbone) on the ssDNA dataset. Each sequence is generated using a flexible backbone representation with either Gaussian noise (b: standard deviation=0.02 Å; c: standard deviation=0.04 Å) or without noise (a: standard deviation=0.00 Å).

**Figure S7:**
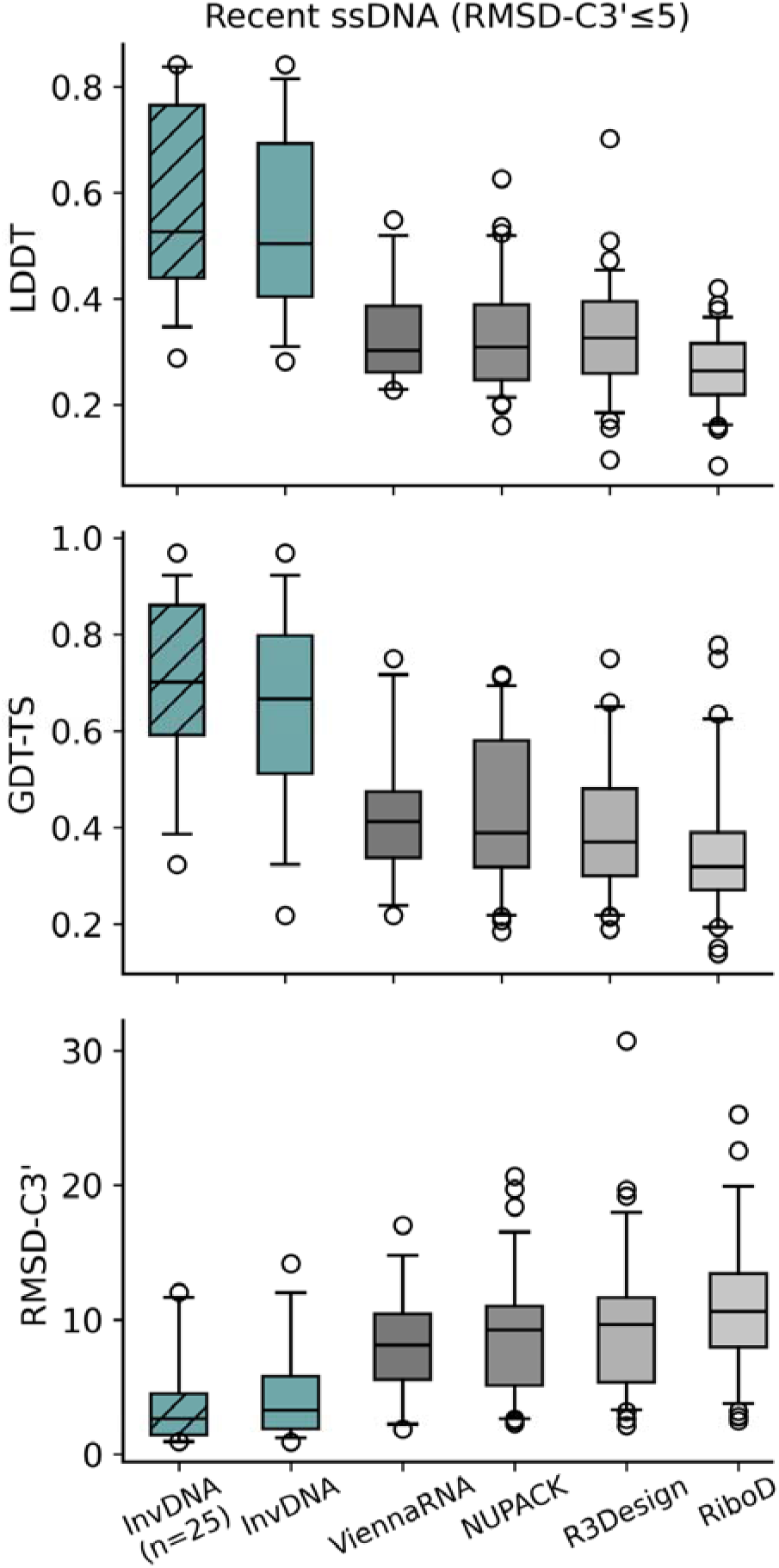
Distribution of LDDT, GDT-TS, and RMSD-C3’ scores for structures predicted by AlphaFold3 on the subset of the recent ssDNA dataset. Predictions were performed on the recent ssDNA dataset using sequences generated by InvDNA, ViennaRNA, NUPACK, R3Design, and RiboDiffusion. For InvDNA (n=25), 25 sequences were generated per ssDNA backbone using a flexible backbone representation with Gaussian noise (standard deviation = 0.04 Å); the sequence yielding the predicted structure with the highest confidence was selected as representative.

**Figure S8:**
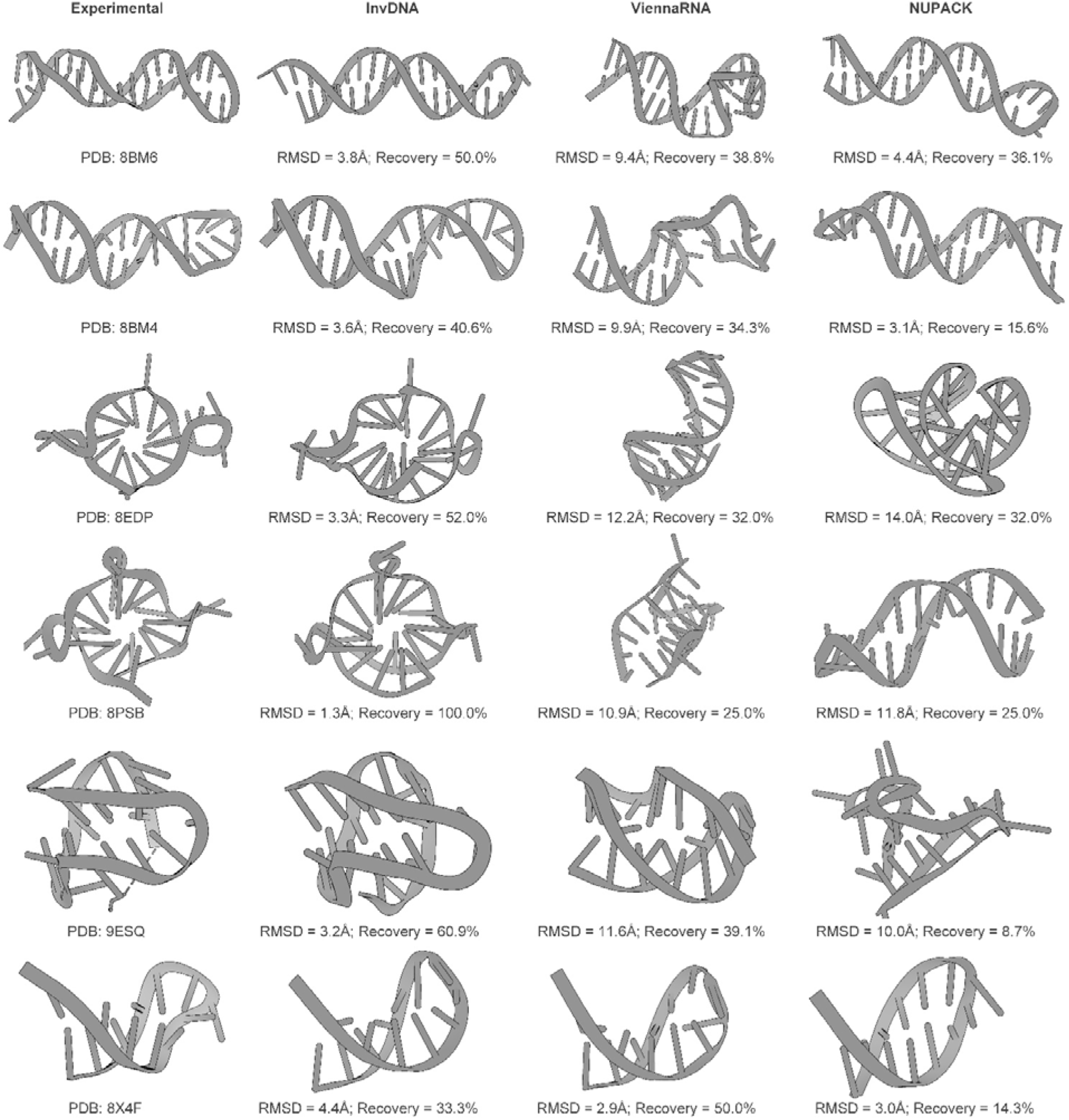
Examples of structures predicted by AlphaFold3 from sequences designed using InvDNA, ViennaRNA and NUPACK. In each row, the left panel shows the target backbone used as the template for sequence design, and all predicted structures are displayed in orientations aligned with their corresponding backbones. All sequences generated by three methods and all structures predicted by AlphaFold3 are available in the supplementary materials.

**Figure S9:**
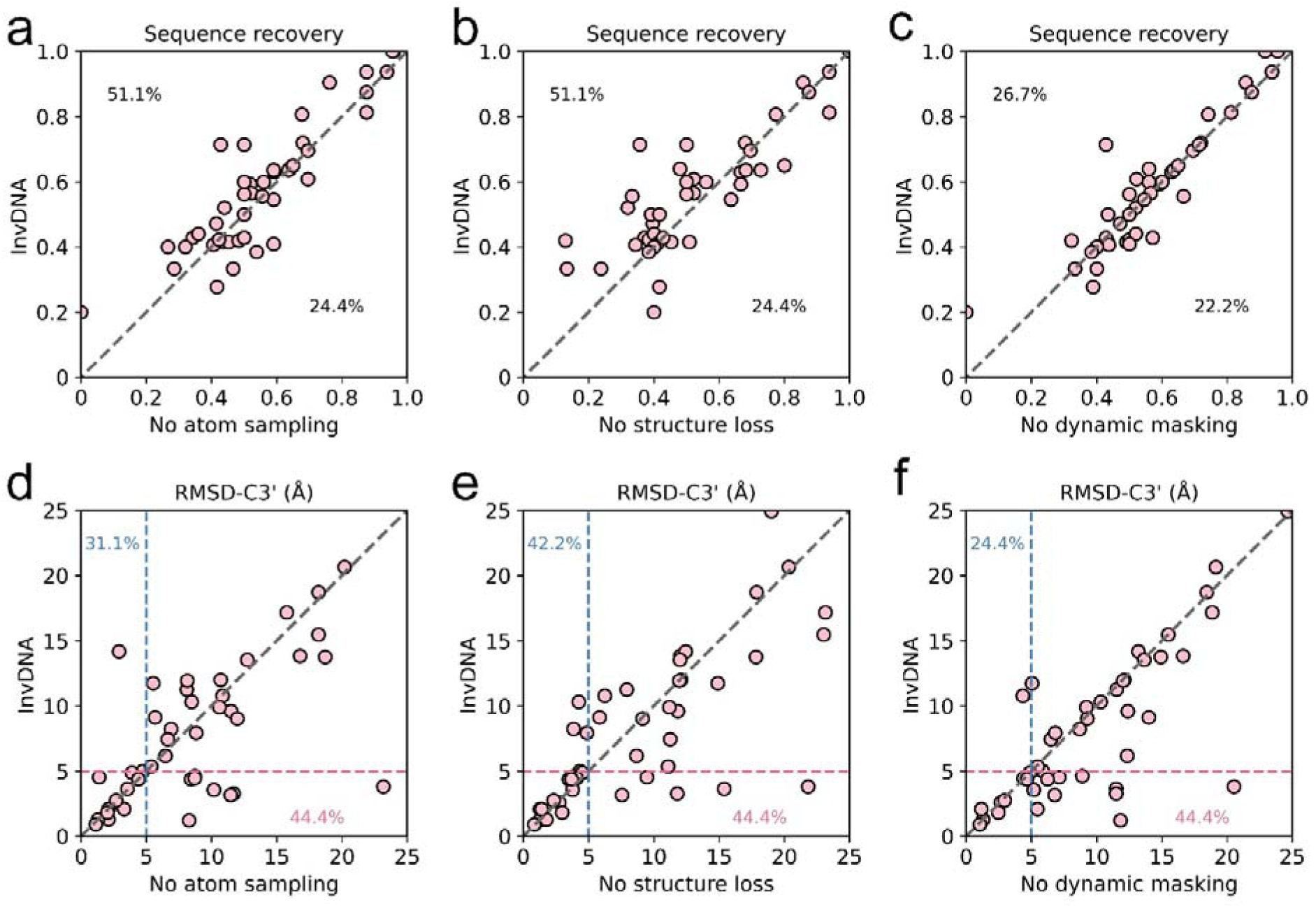
**a-c**, Pairwise comparisons between sequence generated by InvDNA (y-axis, Training data: 100%) and those generated by ablated model variants: **a**, “No atom sampling” model; **b**, “No structure loss” model; **c**, “No dynamic masking” model, on the ssDNA dataset. **d-f**, Pairwise comparisons between structures predicted using sequences generated by InvDNA (y-axis, Training data: 100%) and those generated by ablated model variants: **d**, “No atom sampling” model; **e**, “No structure loss” model; **f**, “No dynamic masking” model, on the ssDNA dataset. The structure is evaluated with RMSD-C3’.

**Figure S10:**
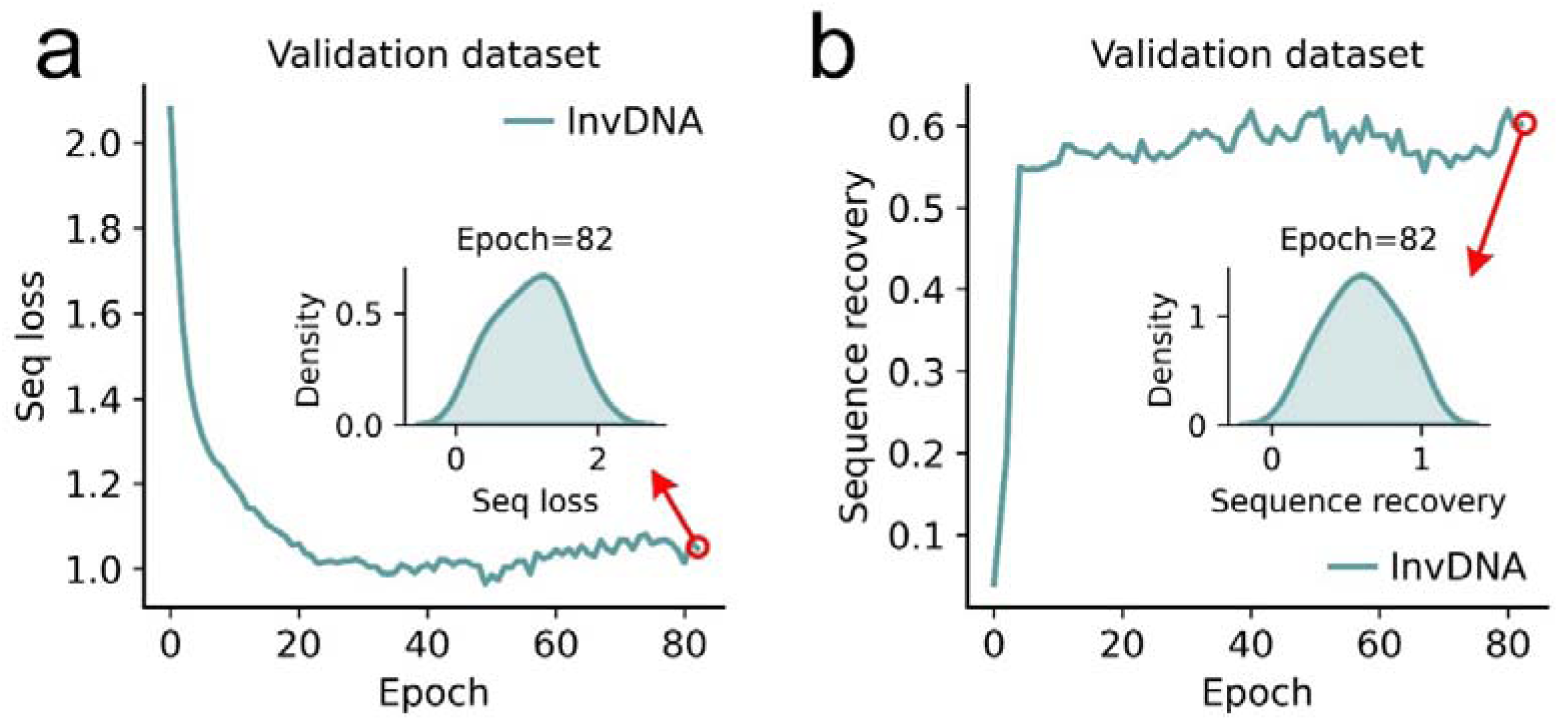
Training curves for InvDNA on the validation dataset: **a:** Mean sequence cross-entropy loss; **d:** Mean sequence recovery rate.

**Figure S11:**
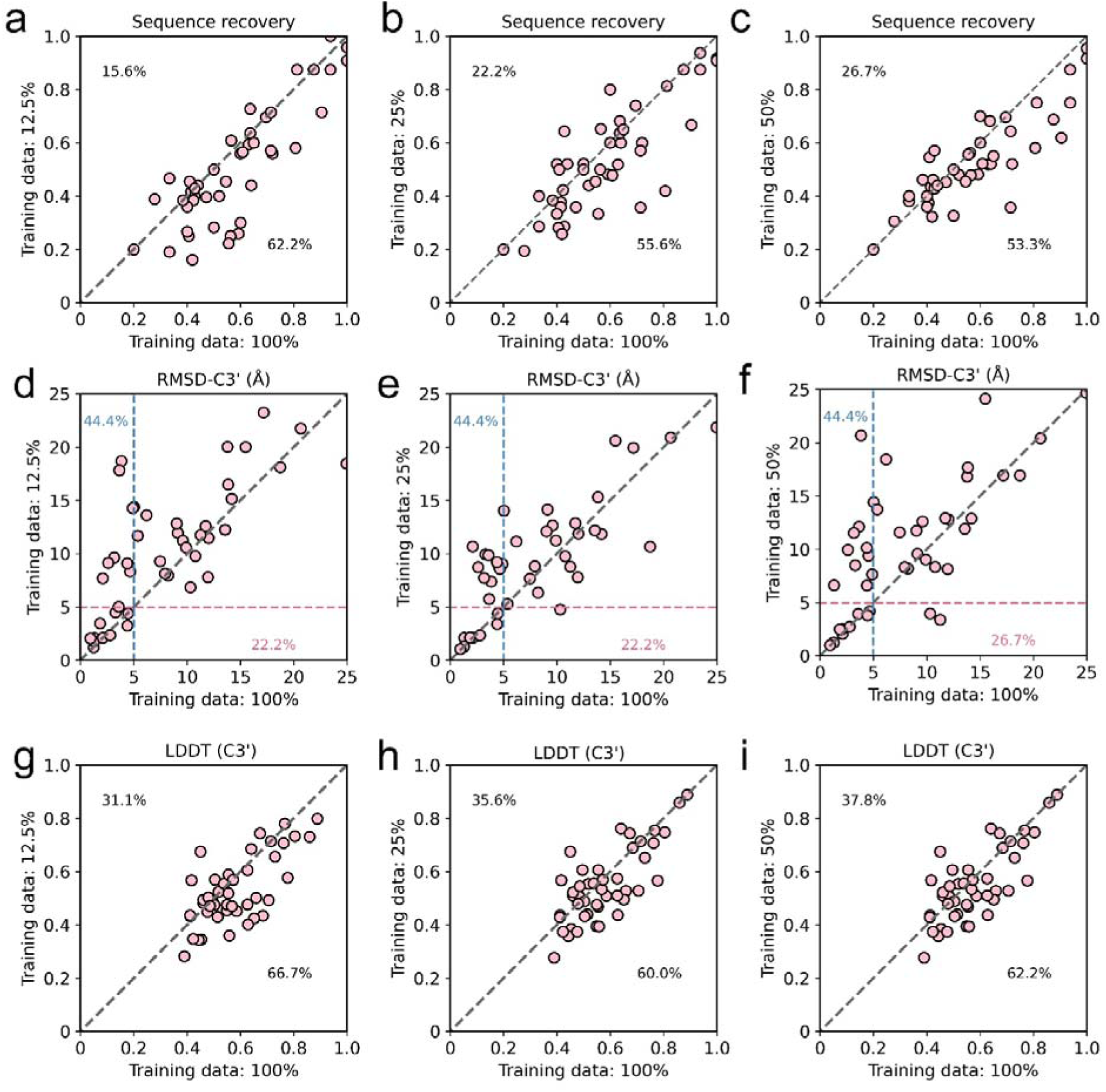
**a-c**, Pairwise comparisons between recovery rate of sequences generated by InvDNA (y-axis, Training data: 100%) and those generated by ablated model variants: **a**, Training data: 12.5%; **b**, Training data: 25%; **c**, Training data: 50%, on the ssDNA dataset. **d-f**, Pairwise comparisons between structures predicted using sequences generated by InvDNA (y-axis, Training data: 100%) and those generated by ablated model variants: **d**, Training data: 12.5%; **e**, Training data: 25%; **f**, Training data: 50%, on the ssDNA dataset. The structure is evaluated with RMSD-C3’. **g-i**, Pairwise comparisons between structures predicted using sequences generated by InvDNA (y-axis, Training data: 100%) and those generated by ablated model variants: **g**, training data: 12.5%; **h**, training data: 25%; **i**, training data: 50%, on the ssDNA dataset. The structure is evaluated with LDDT of C3’ atoms.

## Tables

**Table S1:**
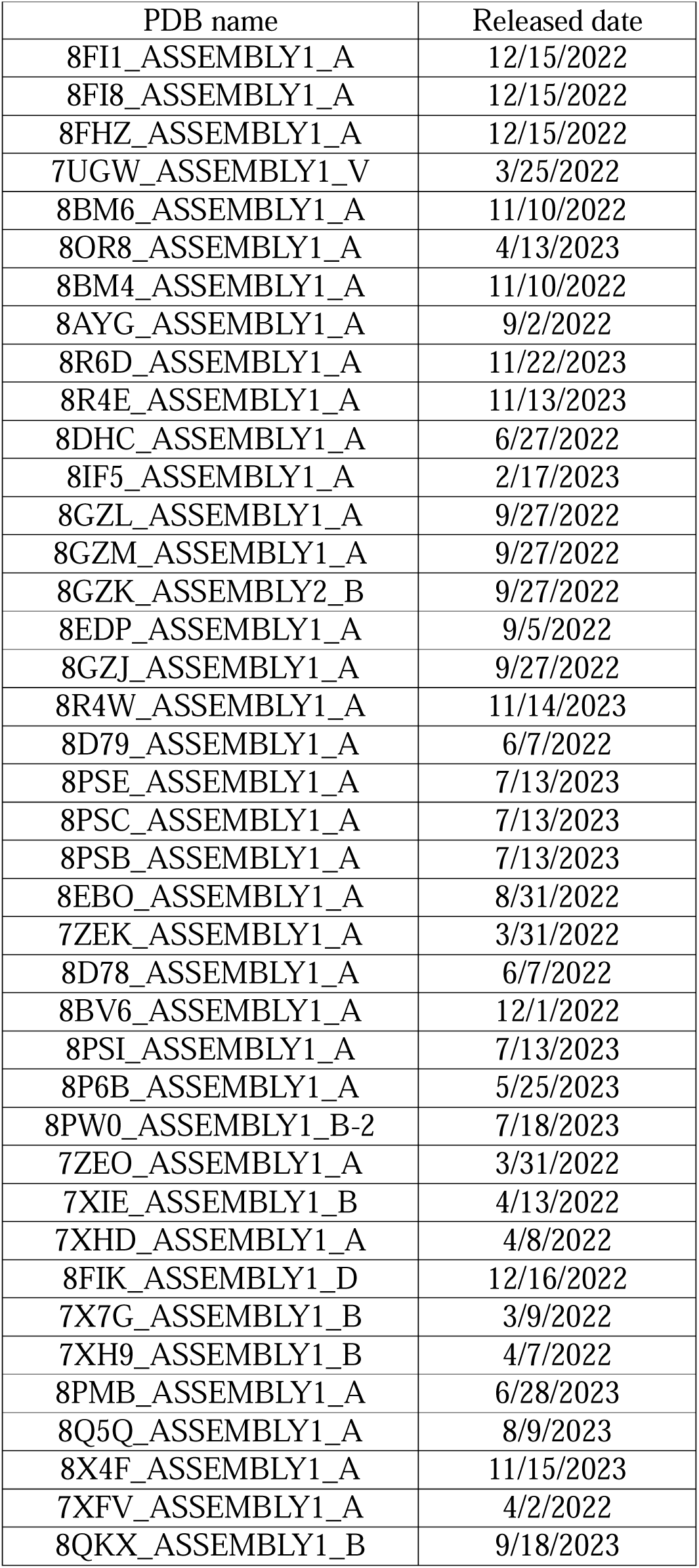
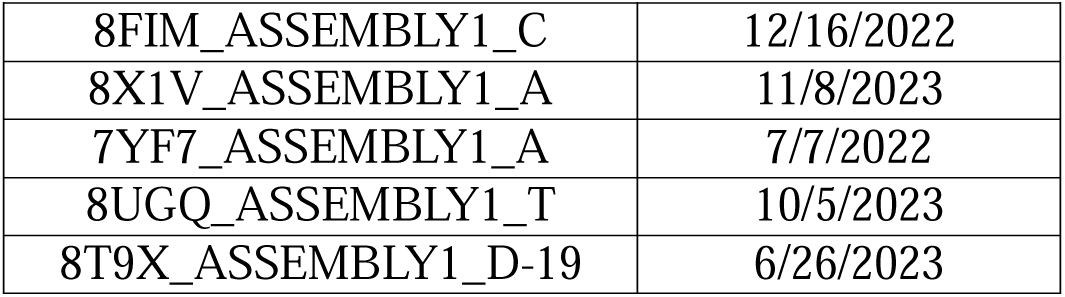
The release date for each ssDNA in the recent ssDNA dataset.

**Table S2:**
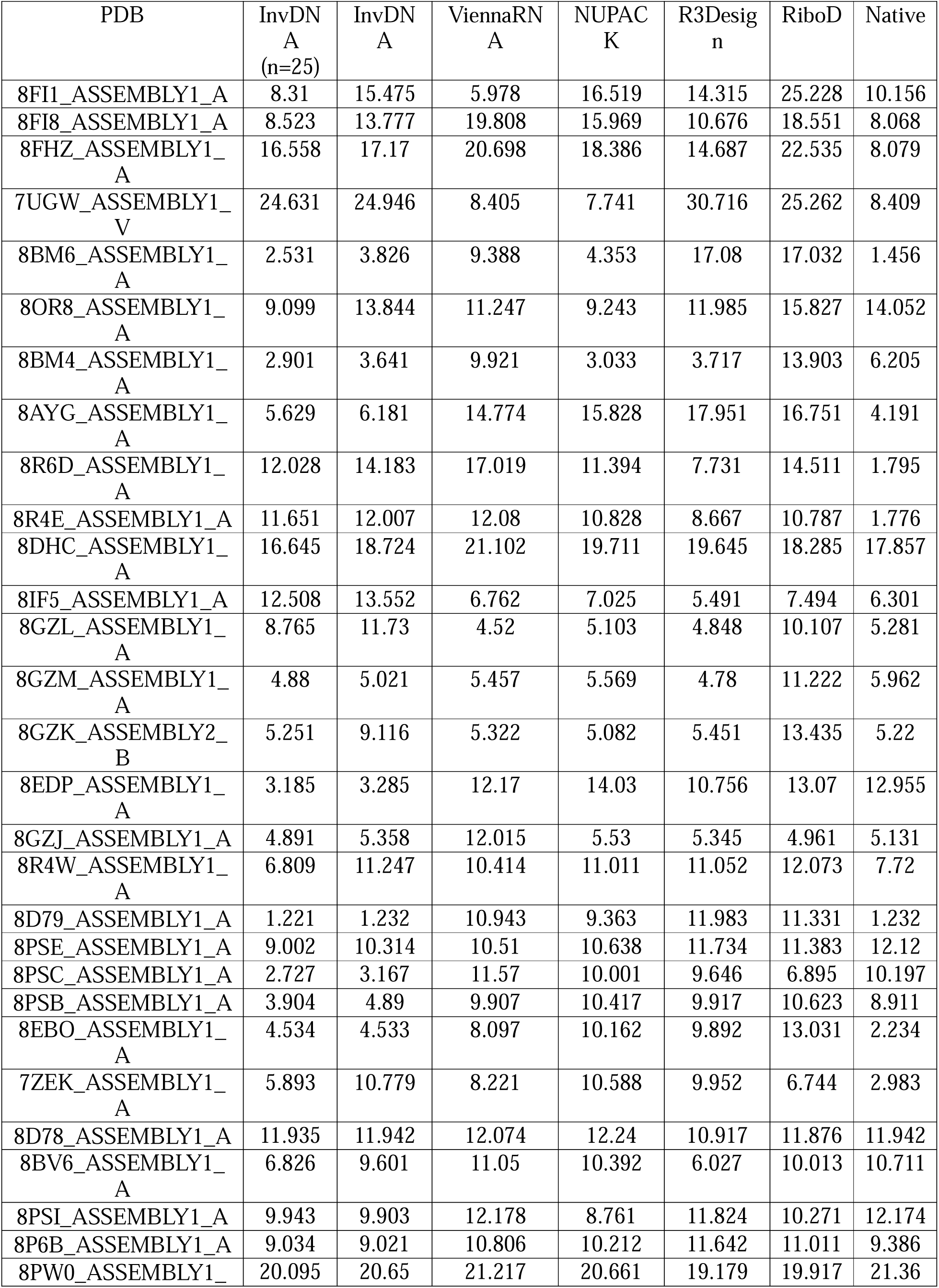
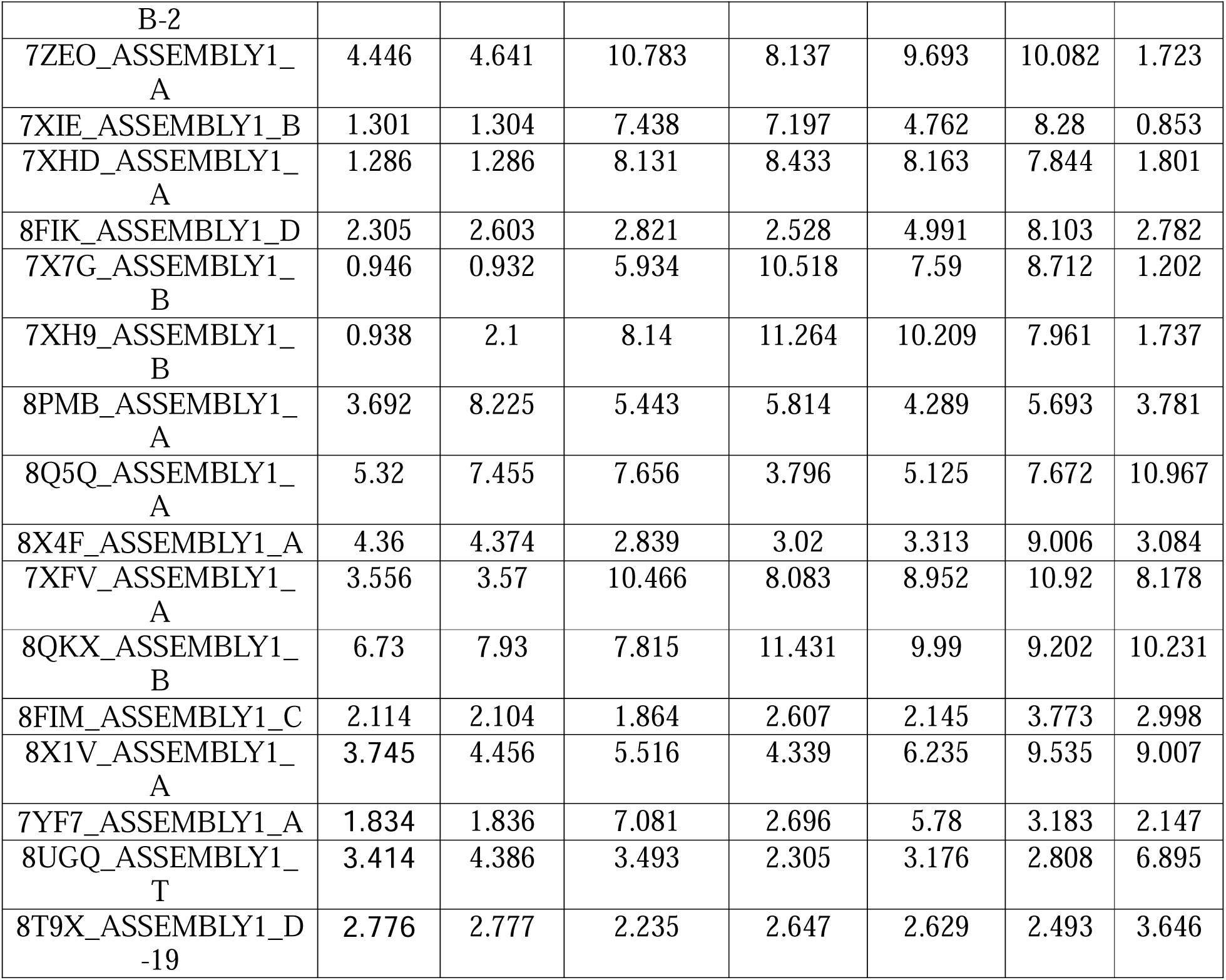
The RMSD-C3’ of structures predicted by AlphaFold3 using sequences generated by InvDNA, ViennaRNA, NUPACK, R3Design, RiboD on the recent ssDNA dataset.

**Table S3:**
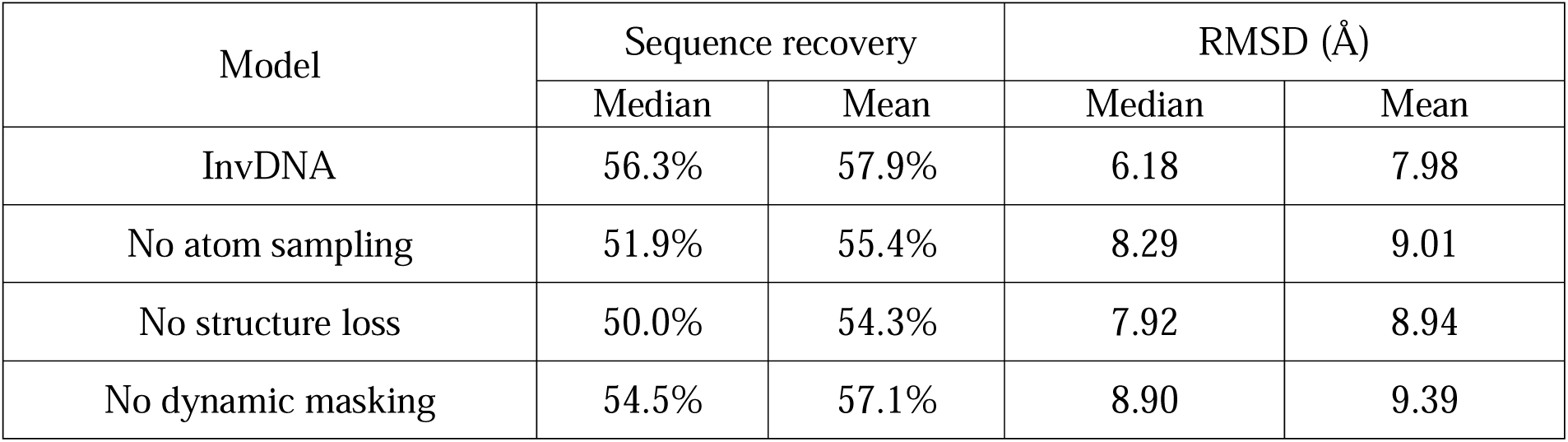
Performance of ablation models used in “Interpreting the InvDNA” on the ssDNA dataset.

**Table S4:** Performance of ablation models used in “Training data size analysis” on the ssDNA dataset.

| Training data | Sequence recovery |  | RMSD (Å) |  | LDDT |  |
| --- | --- | --- | --- | --- | --- | --- |
|  | Median | Mean | Median | Mean | Median | Mean |
| 12.5% | 45.4% | 50.5% | 9.77 | 10.5 | 0.493 | 0.527 |
| 25% | 50.0% | 53.0% | 9.03 | 9.72 | 0.528 | 0.547 |
| 50% | 51.8% | 53.2% | 9.57 | 10.2 | 0.514 | 0.547 |
| 100% | 56.3% | 57.9% | 6.18 | 7.98 | 0.554 | 0.578 |

## References

1. Tian, Z. et al. Circular single-stranded DNA as a programmable vector for gene regulation in cell-free protein expression systems. Nat Commun 15, 4635 (2024).

2. Gozzi, K., Salinas, R., Nguyen, V. D., Laub, M. T. & Schumacher, M. A. ssDNA is an allosteric regulator of the C. crescentus SOS-independent DNA damage response transcription activator, DriD. Genes Dev 36, 618–633 (2022).

3. Hollenstein, M. DNA Catalysis: The Chemical Repertoire of DNAzymes. Molecules 20, 20777–20804 (2015).

4. Chinnapen, D. J.-F. & Sen, D. A deoxyribozyme that harnesses light to repair thymine dimers in DNA. Proceedings of the National Academy of Sciences 101, 65–69 (2004).

5. Scharner, J. & Aznarez, I. Clinical Applications of Single-Stranded Oligonucleotides: Current Landscape of Approved and In-Development Therapeutics. Molecular Therapy 29, 540–554 (2021).

6. Nimjee, S. M., White, R. R., Becker, R. C. & Sullenger, B. A. Aptamers as Therapeutics. Annu Rev Pharmacol Toxicol 57, 61–79 (2017).

7. Malicki, S. et al. Development of selective ssDNA micro-probe for PD1 detection as a novel strategy for cancer imaging. Sci Rep 14, 28652 (2024).

8. Raveendran, M., Lee, A. J., Sharma, R., Wälti, C. & Actis, P. Rational design of DNA nanostructures for single molecule biosensing. Nat Commun 11, 4384 (2020).

9. Sequeira-Antunes, B. & Ferreira, H. A. Nucleic Acid Aptamer-Based Biosensors: A Review. Biomedicines 11, 3201 (2023).

10. Feldkamp, U. & Niemeyer, C. M. Rational design of DNA nanoarchitectures. Angew Chem Int Ed Engl 45, 1856–1876 (2006).

11. Ashrafuzzaman, M., Tseng, C.-Y., Kapty, J., Mercer, J. R. & Tuszynski, J. A. A Computationally Designed DNA Aptamer Template with Specific Binding to Phosphatidylserine. Nucleic Acid Ther 23, 418–426 (2013).

12. Lorenz, R. et al. ViennaRNA Package 2.0. Algorithms for Molecular Biology 6, 26 (2011).

13. Hofacker, I. L. et al. Fast folding and comparison of RNA secondary structures. Monatsh Chem 125, 167–188 (1994).

14. Zadeh, J. N. et al. NUPACK: Analysis and design of nucleic acid systems. J Comput Chem 32, 170–173 (2011).

15. Zadeh, J. N., Wolfe, B. R. & Pierce, N. A. Nucleic acid sequence design via efficient ensemble defect optimization. Journal of Computational Chemistry 32, 439–452 (2011).

16. Vicens, Q. & Kieft, J. S. Thoughts on how to think (and talk) about RNA structure. Proceedings of the National Academy of Sciences 119, e2112677119 (2022).

17. Abraham, M., Dror, O., Nussinov, R. & Wolfson, H. J. Analysis and classification of RNA tertiary structures. RNA 14, 2274–2289 (2008).

18. Improved free-energy parameters for predictions of RNA duplex stability. https://www.pnas.org/doi/epdf/10.1073/pnas.83.24.9373doi:10.1073/pnas.83.24.9373.

19. Jaeger, J. A., Turner, D. H. & Zuker, M. Improved predictions of secondary structures for RNA. Proc Natl Acad Sci U S A 86, 7706–7710 (1989).

20. Xia, T. et al. Thermodynamic parameters for an expanded nearest-neighbor model for formation of RNA duplexes with Watson-Crick base pairs. Biochemistry 37, 14719–14735 (1998).

21. Tan, C. et al. R3Design: deep tertiary structure-based RNA sequence design and beyond. Brief Bioinform 26, bbae682 (2025).

22. Dauparas, J. et al. Robust deep learning–based protein sequence design using ProteinMPNN. Science 378, 49–56 (2022).

23. Liu, Y. et al. Rotamer-free protein sequence design based on deep learning and self-consistency. Nat Comput Sci 2, 451–462 (2022).

24. Joshi, C. K. et al. gRNAde: Geometric Deep Learning for 3D RNA inverse design. bioRxiv 2024.03.31.587283 (2025) doi:10.1101/2024.03.31.587283.

25. Anand, N. et al. Protein sequence design with a learned potential. Nat Commun 13, 746 (2022).

26. Zhang, Y., Xiong, Y. & Xiao, Y. 3dDNA: A Computational Method of Building DNA 3D Structures. Molecules 27, 5936 (2022).

27. Wang, J., Wang, J., Huang, Y. & Xiao, Y. 3dRNA v2.0: An Updated Web Server for RNA 3D Structure Prediction. International Journal of Molecular Sciences 20, 4116 (2019).

28. Hong, X., Zhan, J. & Zhou, Y. On the completeness of existing RNA fragment structures. 2024.05.06.592843 Preprint at 10.1101/2024.05.06.592843 (2024).

29. Ahdritz, G. et al. OpenFold: retraining AlphaFold2 yields new insights into its learning mechanisms and capacity for generalization. Nat Methods 21, 1514–1524 (2024).

30. Ren, M., Yu, C., Bu, D. & Zhang, H. Accurate and robust protein sequence design with CarbonDesign. Nat Mach Intell 6, 536–547 (2024).

31. Dauparas, J. et al. Atomic context-conditioned protein sequence design using LigandMPNN. Nat Methods 22, 717–723 (2025).

32. Jumper, J. et al. Highly accurate protein structure prediction with AlphaFold. Nature 596, 583–589 (2021).

33. Abramson, J. et al. Accurate structure prediction of biomolecular interactions with AlphaFold 3. Nature 630, 493–500 (2024).

34. Huang, H., Lin, Z., He, D., Hong, L. & Li, Y. RiboDiffusion: tertiary structure-based RNA inverse folding with generative diffusion models. Bioinformatics 40, i347–i356 (2024).

35. Berman, H. M. et al. The Protein Data Bank. Nucleic Acids Res 28, 235–242 (2000).

36. Lu, X.-J., Bussemaker, H. J. & Olson, W. K. DSSR: an integrated software tool for dissecting the spatial structure of RNA. Nucleic Acids Res 43, e142 (2015).

37. Tan, C. et al. R3Design: deep tertiary structure-based RNA sequence design and beyond. Brief Bioinform 26, bbae682 (2025).

38. RR3DD: an RNA global structure-based RNA three-dimensional structural classification database.

39. Bu, F. et al. RNA-Puzzles Round V: blind predictions of 23 RNA structures. Nat Methods 22, 399–411 (2025).

40. Svozil, D., Kalina, J., Omelka, M. & Schneider, B. DNA conformations and their sequence preferences. Nucleic Acids Res 36, 3690–3706 (2008).

41. McPartlon, M. & Xu, J. An end-to-end deep learning method for protein side-chain packing and inverse folding. Proceedings of the National Academy of Sciences 120, e2216438120 (2023).

42. Huang, X., Pearce, R. & Zhang, Y. FASPR: an open-source tool for fast and accurate protein side-chain packing. Bioinformatics 36, 3758–3765 (2020).

43. Eastman, P. et al. OpenMM 7: Rapid development of high performance algorithms for molecular dynamics. PLOS Computational Biology 13, e1005659 (2017).

44. Zhu, J. et al. StruCloze: A Unified Framework for Backmapping and Inpainting of Biomolecules. 2025.06.26.661889 Preprint at 10.1101/2025.06.26.661889 (2025).

45. Li, W., Jaroszewski, L. & Godzik, A. Clustering of highly homologous sequences to reduce the size of large protein databases. Bioinformatics 17, 282–283 (2001).

46. Kagaya, Y. et al. NuFold: end-to-end approach for RNA tertiary structure prediction with flexible nucleobase center representation. Nat Commun 16, 881 (2025).

47. Biasini, M. et al. OpenStructure: an integrated software framework for computational structural biology. Acta Cryst D 69, 701–709 (2013).

